# Erasable Hippocampal Neural Signatures Predict Memory Discrimination

**DOI:** 10.1101/2023.02.02.526824

**Authors:** Nathaniel R. Kinsky, Daniel J. Orlin, Evan A. Ruesch, Brian Kim, Siria Coello, Kamran Diba, Steve Ramirez

**Author notes:** These authors contributed equally to this work (Nathaniel R. Kinsky, Daniel J. Orlin). Corresponding authors. Correspondence should be addressed to N.R.K or S.R.

## Abstract

Memories involving the hippocampus can take several days to consolidate, challenging efforts to uncover the neuronal signatures underlying this process. Using calcium imaging in freely moving mice, we tracked the hippocampal dynamics underlying memory formation across a ten-day contextual fear conditioning (CFC) task. Following learning, context-specific place field remapping correlated with memory performance. To causally test whether these hippocampal dynamics support memory consolidation, we induced amnesia in a group of mice by pharmacologically blocking protein synthesis immediately following learning. We found that halting protein synthesis following learning paradoxically accelerated cell turnover and also arrested learning-related remapping, paralleling the absence of remapping observed in untreated mice that exhibited poor memory expression. Finally, coordinated neural activity that emerged following learning was dependent on intact protein synthesis and predicted memory-related freezing behavior. We conclude that context-specific place field remapping and the development of coordinated ensemble activity require protein synthesis and underlie contextual fear memory consolidation.

## Introduction

The process of memory consolidation requires protein synthesis following learning to produce stable long-term memories (Barondes & Cohen, 1968; Davis & Squire, 1984; Squire & Barondes, 1974). At the synaptic level, protein synthesis is necessary for the formation of late-phase but not early-phase long-term potentiation (LTP) in hippocampal (HPC) neurons (Frey & Morris, 1997; Huang et al., 1994; Nguyen et al., 1994). At the behavioral level, blocking protein synthesis following learning impairs long-term memory for hippocampus-dependent contextual fear conditioning (CFC) while leaving short term memory intact (Ryan et al., 2015; Schafe et al., 1999). These studies suggest that memory consolidation requires new proteins to reinforce learning-related connections which are potentiated during learning. Despite the evidence linking *in vitro* synaptic potentiation to memory consolidation, little is known about how protein synthesis *in vivo* influences the functional coding properties of neurons before, during, and after learning to support memory consolidation. Experience-dependent remapping of place fields in the hippocampus, which requires NMDA receptor dependent-plasticity (Dupret et al., 2010), is thought to reflect learning-related reorganization of the hippocampal network (Bostock et al., 1991; Colgin et al., 2008). Therefore, to causally test the hypothesis that lasting, learning-related place-cell remapping (Muller & Kubie, 1987) is necessary for consolidation of a CFC memory (Moita et al., 2004; Wang et al., 2012) we combined *in vivo* calcium imaging in mice with systemic administration of the protein synthesis inhibitor, anisomycin, and tracked the evolution, remapping, and stabilization of HPC place fields during and after CFC. Finally, we leveraged the ability to record from large cell ensembles to investigate whether blocking protein synthesis disrupted the development of coordinated neural activity predictive of freezing behavior. Our results indicate that context-specific remapping and coordinated, freeze-predicting ensemble activity emerge in a protein-synthesis dependent manner following learning to support CFC memory consolidation.

## Results

### Hippocampal neural dynamics between arenas predicts memory specificity

Weeks prior to training, all mice received viral infusions of the genetically encoded calcium indicator GCaMP6f (Chen et al., 2013) in region CA1 of the dorsal hippocampus. Subsequently, following two days of pre-exposure (“Before” learning, days −2 and −1) to an open field (neutral arena) and an operant chamber (shock arena) mice received a mild foot-shock on day 0 (training), after which they were moved to their home cage and immediately given systemic injections of either anisomycin (ANI group) or vehicle (CTRL group). The shock arena recording immediately followed each neutral arena recording, separated by approximate 5 minutes to disconnect the recording camera and move the mouse the other room/arena. We then performed a test of short-term memory 4 hours later and three tests of long-term memory 1, 2, and 7 days following training (Figure 1A), each time by measuring freezing behavior. We titrated the shock level during training such that the mice froze significantly more in the shock arena relative to the neutral arena following learning while still exploring the majority of both arenas. Nonetheless, we observed a range of freezing levels during the day 1 and 2 memory tests (Figure S1A) and subsequently sub-divided CTRL mice into two groups: Learners, who froze significantly more in the shock arena, and Non-Learners, who either showed generalized freezing or froze at low levels in both arenas (Figures 1B and Figure S1B,E). Learners exhibited reduced freezing on day 7, indicative of extinction (Figure 1B). All subsequent analyses related to long-term memory therefore utilized the day 1 and 2 recall sessions (“After” learning and consolidation). In contrast, mice in the ANI group exhibited no difference in freezing between arenas at any time point, indicating that blocking protein synthesis impaired context-specific fear memory (Figure 1C). To assess memory specificity, we calculated a behavioral discrimination index (DI_beh_) which quantified how much each animal froze in the shock vs. neutral arenas. Negative DI values indicated higher freezing in the shock arena. Learners exhibited negative DI_beh_ levels on days 1 and 2 that were significantly different from both Non-Learners (by definition) and from the ANI group as well (Figure 1D). Variability in freezing on the days prior to shock could indicate heightened anxiety by some mice, which could in turn influence contextual fear learning. However, we found no difference between groups in thigmotaxis, a metric of anxiety (Figure S1F-G). Both the CTRL and ANI groups exhibited significant freezing in the shock arena during the 4 hour test, though we note that the behavior in the ANI group at this time point could arise from either contextual fear or non-specific aversive effects of anisomycin treatment (Figure S2) since these mice froze at high levels in both arenas.

**Figure 1:**
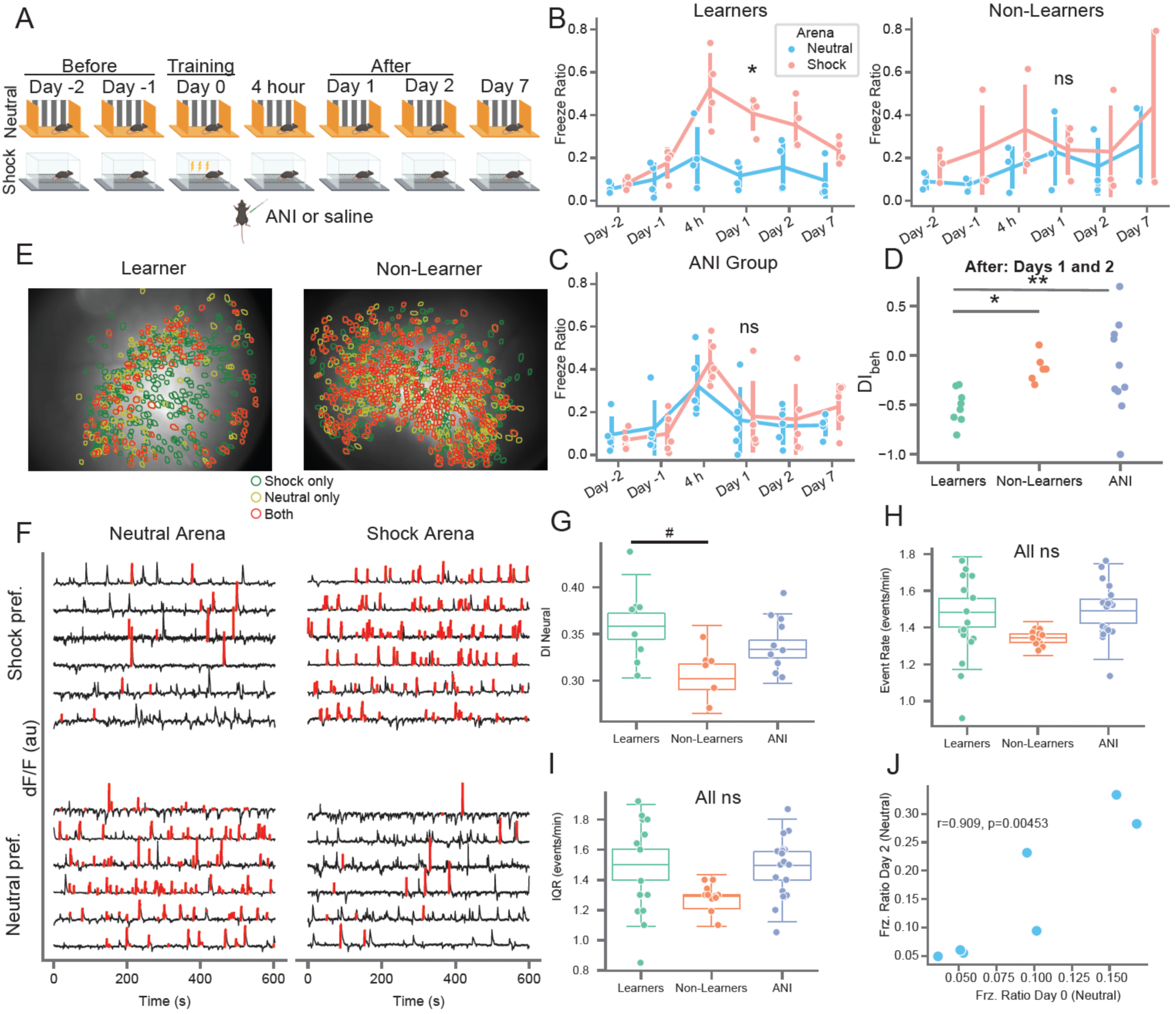
Mice exhibit variability in memory recall and neural activity prior to learning in a contextual fear conditioning task . **A)** Schematic of the behavioral paradigm. Mice freely explored two distinct arenas (neutral and shock) for 10 minutes each day. Mice underwent mild contextual fear conditioning on day 0 in the shock arena followed by immediate I.P. administration of anisomycin or vehicle in their home cage. Memory recall tests were conducted 4 hours and 1, 2, and 7 days post-shock. The time of each session is referenced to the shock session. **B)** (left) Learner (CTRL) mice freezing on all days. Red = shock arena, blue = neutral arena. *p=4.5e-0.4 shock – neutral freezing from day-1 to day 1 one-sided paired t-test (n=4 mice, t=13.4). (right) Same but for Non-Learner (CTRL) mice (n=3 mice, p=0.249, t=0.819). **C)** Same as B but for ANI group (n=5 mice, p=0.219, t=0.859). **D)** Behavioral discrimination between arenas after shock (Days 1-2) shows formation of a specific fear memory for Learners only, by definition (positive = more freezing in neutral arena, negative = more freezing in shock arena, 0 = equal freezing in both arenas). *p=0.009 (t=3.56), #p=0.06 (t=1.83) 1-sided t-test of mean DI value from Days 1 & 2, n=8/6/10 sessions for Learners/Non-Learners/ANI group. **E)** (left) Neural overlap plots between neutral and shock arenas for an example Learner mouse on day −1, before shock. Green = cells active in the shock arena only, yellow = cells active in the neutral arena only, orange = cells active in both arenas. (right) Same for example Non-Learner on day −2 showing higher overlap of active cells between arenas. **F)** Example calcium activity from the Learner mouse shown in C (left) for cells active in both arenas. Black = calcium trace, red = putative spiking activity during transient rises. Top row shows shock arena preferring cells, bottom row shows neutral arena preferring cells. **G)** Neural discrimination index (DIneural) between groups on Days −2 and −1. Boxplots show population median and 1^st^/3^rd^ quartiles (whiskers, 95% CI) estimated using hierarchical bootstrapping (HB) data with session means overlaid in dots, #p=0.09 after Bonferroni correction for multiple comparisons**. H)** Same as G) but for event rate in Shock arena, **I**) Same as G) but for event rate interquartile range (IQR) in Shock arena. **J)** Freezing in Neutral arena on Day 2 vs. Neutral arena on Day 0. Pearson correlation value and p-value (two-sided) shown on plot. Statistics for G-J: un-paired one-sided HB test for days −2 and −1 after Bonferroni correction, n=10,000 bootstraps.

We visualized the activity of pyramidal neurons in region CA1 of the dorsal hippocampus (Figure 1E) using a miniaturized epifluorescence microscope (Ghosh et al., 2011; Ziv et al., 2013). We identified a large number of neurons in each 10 minute session (n = 128 to 1216, Figure S3), extracted their corresponding calcium traces (Figure 1F), and tracked them between sessions throughout the CFC task, which allowed us to determine the long-term evolution of the HPC neural code. We noticed that many neurons exhibited strong changes in mean event rate between arenas (Figure 1F) and calculated a neural discrimination index (DI_neural_) to quantify the distinctiveness of neural activity between arenas (0 = similar, 1 = distinct). Same-day neural discrimination between arenas correlated strongly with across-day neural discrimination in the same arena (Figure S1D). This indicates that mice exhibit natural variability in neural discrimination which is invariant between different arenas and across time.

Surprisingly, we noticed that Learners exhibited a trend toward higher DI_neural_ values compared to Non-Learners in the sessions prior to the shock that did not reach significance (Figure 1G). This was not due to higher activity rates for Non-Learners since we found no differences between groups in mean event rate and or distribution of event rates on the days prior to shock (Figure 1H-I). Finally, we found that freezing behavior the day of training correlated with Neutral arena freezing during the 4 Hour and Day 2 recall sessions and exhibited a trend toward correlation on Days 1 and 7 (Figure 1J and Figure S1H). These results indicate that the animal’s behavioral state the day of conditioning, and potentially neural discrimination between arenas prior to learning, can influence memory specificity.

### Blocking protein synthesis disrupts persistent neural activity following learning and arrests learning-related place field remapping

Next, we probed how arresting protein synthesis impacted HPC dynamics. Previous studies by us and others have found that hippocampal cells exhibit constant turnover over time such that the subset of active cells slowly changes over time (Kinsky et al., 2018; Ziv et al., 2013). We hypothesized that, by preventing plasticity, ANI administration would slow or stop the normal rate of cell turnover, measured as the overlap of active cells at each timepoint with the first day recording day (Figure 2A). Surprisingly, ANI administration temporarily accelerated cell turnover between the Day −1 and 4 hour session after which turnover rate returned to normal (Figure 2B, C). We noticed that the normalized number of active neurons also appeared to be lower for the ANI group compared to the CTRL group at the 4 hour session when anisomycin was still on board, and potentially the following sessions, which could contribute to increased turnover (Figure 2D). However, diminished cell activity could also result from reduced locomotion (Rich et al., 2014), which we observed in both groups following conditioning / anisomycin. To disentangle the effects of anisomycin and locomotion, we therefore fit the normalized number of cells observed in each recording session to a generalized linear model with arena, freezing ratio, anisomycin status (acute = 4 hr session, after = Day 1, 2, and 7), experimental group, and anisomycin status x experimental group as covariates (Methods). Under this framework, we found a highly significant influence of freezing ratio on active cell number (p=2.3x10^-5^) and a trend toward fewer active cells for the ANI group at the 4 hour session (p=0.056) and following sessions (p=0.094) that did not reach significance (Figure 2D). These results indicate that reduced locomotion is the primary driver of reduced cell activity observed following anisomycin administration. To test whether reduced activity resulted from cell death, we injected a separate cohort of mice with either saline or anisomycin and then performed immunohistochemistry to stain for apoptosis 4 hours after injection. We found negligible levels of apoptosis overall with no difference between groups (Figure S4). We also observed no difference in the mean height of calcium transients (a measure of signal-to-noise ratio) for neurons active before, during, or after ANI administration, indicating that the observed decrease in the number of active neurons was not due to differences in Ca2+ levels or depletion of the GCaMP protein (Figure S5D-E). To confirm that anisomycin administration did not suppress network activities, which was reported for intracranial infusions (but not systemic injections) of protein synthesis inhibitors in anesthetized or immobile rodents (Barondes & Cohen, 1966; Sharma et al., 2012: Park et al., 2023), we performed extracellular recordings in freely moving rats implanted with a linear probe in the CA1 region of the hippocampus following administration of anisomycin. We observed preserved theta activity, theta modulation of spiking, and robust sharp wave ripple activity for up to 5 hours following anisomycin administration (Figure S5A-C). These observations indicate that blocking protein synthesis following learning does not simply abolish all neuronal activity but instead acutely accelerates the turnover of cells which may be specifically involved in learning.

**Figure 2:**
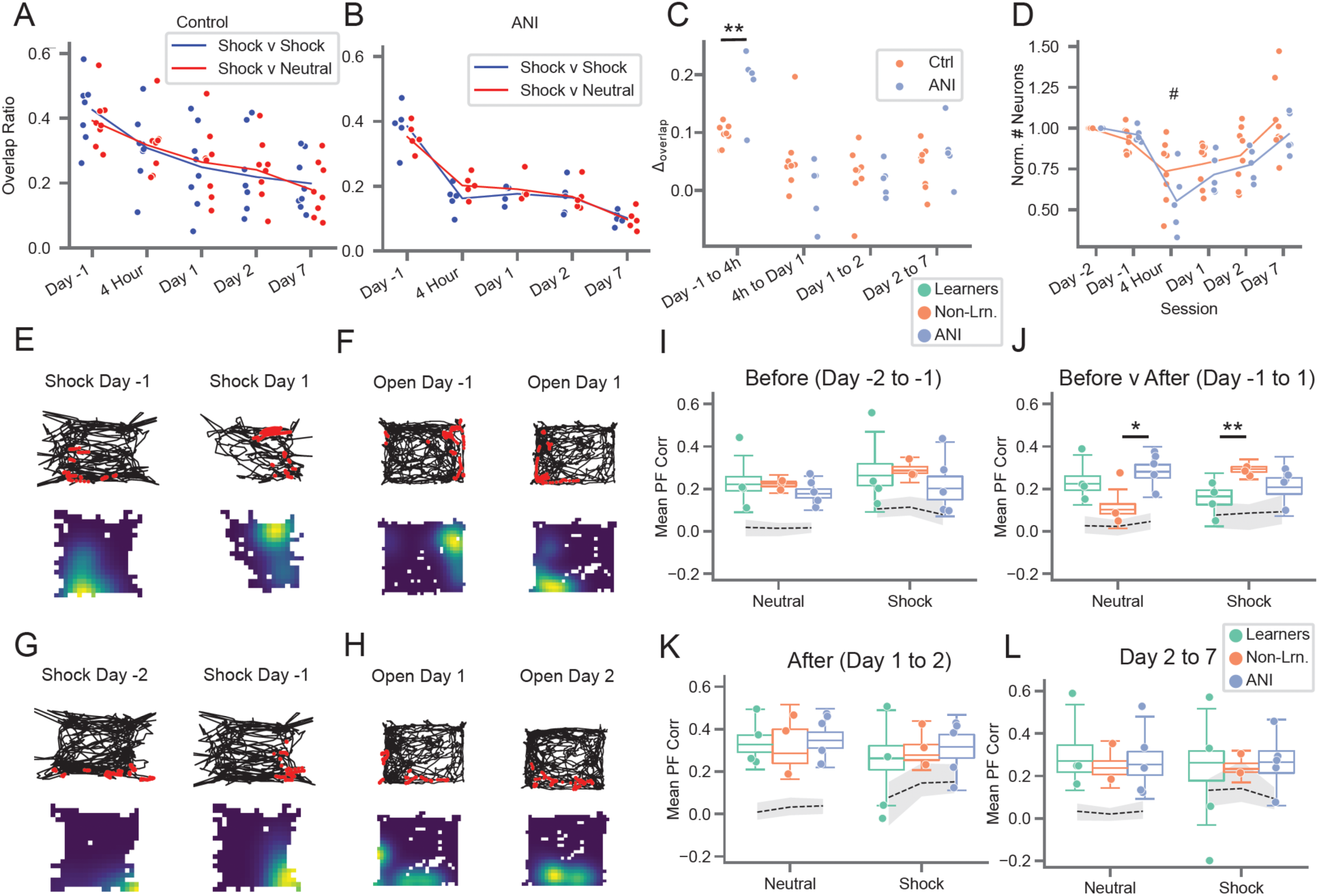
Preventing protein synthesis accelerates cell turnover and stifles learning-related place field remapping. **A)** Cell overlap ratio with Day −2 session, CTRL group. Blue = within shock arena, red = shock v. neutral arena. **B)** Same as A) but for ANI group. **C)** Change in overlap ratios from A) and B), dots show values from both arenas for each mouse, **p=0.00174 two-sided t-test of mean value for each mouse (t=4.11, n=7 CTRL mice and 5 ANI mice). **D)** Number of active neurons observed each day, normalized to day −1. p=2.e-5 freeze-ratio, #p=0.056 group x 4 hr session interaction, p=0.094 group x after interaction, generalized linear model. **E)** and **F)** Example place fields exhibiting learning-related remapping. **E)** Place field in shock arena from Learner mouse. (*top)* Example mouse trajectory (black) with calcium activity (red) overlaid for the same cell from day −2 to −1 in shock arena, *(bottom)* occupancy normalized rate maps for the same cells with warm colors indicating areas of high calcium activity. **F)** Same as E) but for Non-Learner mouse in Neutral arena. **G)** and **H)** Example stable place fields. **G)** Same as E) but for a different cell from same mouse in the shock arena prior to conditioning. **H)** Same as F) but for a different cell from the same mouse in the neutral arena after conditioning. **I)** Place field correlations for all mice before shock (Days −2 and −1), boxplots show population median and 1^st^/3^rd^ quartiles (whiskers, 95% CI) estimated using hierarchical bootstrapping (HB) data with session means overlaid in dots. Dashed line and grey shading show mean and 95% CI of correlations calculated from shuffling cell identify 1000 times between sessions. **J)** Same as I) but for Day −1 to Day 1, *p=0.0496, **p=0.0034. **K)** Same as I) but for Day 1 to Day 2. **L)** Same as I) but for Day 2 to Day 7. Statistics for I-L: un-paired one-sided HB test after Bonferroni correction, n=10,000 bootstraps.

We next hypothesized that arresting protein synthesis, which disrupts the permanence of newly formed place fields (Agnihotri et al., 2004), would likewise impair the long-term stability of CFC-related remapping (Moita et al., 2004; Wang et al., 2012). We therefore assessed place field remapping within and across epochs by correlating event rate maps for all neurons active between two sessions (Figure 2E-L) following administration of anisomycin, compared with saline. We noticed that place fields recorded in the Learners group exhibited very low across-session correlations in the neutral arena throughout the experiment (Figure S6), which could indicate remapping. However, in a separate study, we observed what appeared to be remapping between two recording sessions, but which was actually a coherent rotation of all place fields around a singular point (Kinsky et al., 2018), indicative of confusion in an animal’s axis of orientation, such as between west and north (Keinath et al., 2017). Importantly, such angular disorientation does not impair the ability of mice to identify specific places (Julian et al., 2015), and between-cell firing relationships are maintained during coherent rotations whereas they are scrambled following remapping. Coherent rotations match the geometry of the environment, e.g. producing rotations in 60 degree increments in a triangle versus 90 degrees in square environments such as our neutral and shock arenas. Therefore, to ensure we were not mistaking coherent rotations for remapping, we rotated all place field maps from each session in 90 degree increments and used only the orientation that produced the highest correlation between sessions.

We found generally higher correlations compared to using non-rotated maps (Figure 2I-L vs. Figure S6F), confirming that coherent rotations occurred in many recording sessions. In particular, place field correlations were high both before learning and after consolidation for all groups (Figure 2I, K, L and Figure S7), though Learner correlations trended toward lower stability after shock (Figure 2K, L) which could indicate extinction (Wang et al., 2015). We then compared place fields between session to assess short-term (Figure S8, 4 hour session to Days −1 and 1) and long-term (Figure 2J, Day −1 to Day 1) learning-related remapping. In agreement with previous studies (Moita et al., 2004; Wang et al., 2012), Learner place fields remapped from Day −1 to Day 1, as indicated by lower correlations in the shock arena compared to the other groups (Figure 2J, Figure S7B). Interestingly, Non-Learners exhibited lower place map correlations from Day −1 to Day 1 in the neutral arena compared to the other groups, indicating paradoxically stable place fields in the shock arena but remapping in the neutral arena (Figure 2J, Figure S7C, Figure S8C). We observed similar results when examining population vector correlations (Figure S9). If learning causes remapping, this double dissociation indicates that Non-Learner memory deficits might result from improperly associating the neutral arena with shock. In contrast, the place maps of mice in the ANI group displayed high correlations in both arenas across all time points, including after learning (Figure 2I-L, Figure S7E-F). This deficit in remapping following anisomycin therefore indicates that protein synthesis is required to stabilize the set of place fields which emerge following learning to support memory consolidation. Overall, these observations indicate that remapping is necessary for the creation of context-specific memories and that a lack of remapping, or improperly remapping, may underlie the memory deficits observed in Non-Learners and in the ANI group.

### Coordinated freeze-predictive neural activity emerges following consolidation

In addition to the spatial code, hippocampal neural activity can also reflect non-spatial variables in a task (McKenzie et al., 2014; Muzzio et al., 2009; Wood et al., 1999). We noticed that, in line with recent studies (Lee & Han, 2022; Schuette et al., 2020), many hippocampal neurons reliably produced calcium transients around the time that mice froze (Figure 3A-B, D-F and S10A-C). We observed similar proportions of peri-freeze tuned cells across all groups and recording sessions, even before CFC (Figure S10D-E). Examining the +/-4 seconds around freezing events for Days 1-2 (after learning) revealed that population-level neural activity peaked in the 2 seconds prior to freezing for control mice compared to the ANI group (Figure 3C) but did not differ between Learners and Non-Learners (Figure S10F). This pre-freeze increase in activity for the CTRL vs. the ANI group was not present before learning or at the 4 hour session (Figure S10L-M). After learning, neurons with significant peri-freeze tuning tiled the entire +/-4 seconds around freeze onset (Figure 3G). However, the average peak timing of cells with significant peri-freeze activity shift shifted to significantly early time points for Learners vs. Non-Learners and ANI mice after learning (Figure 3H) with ∼70% of cells active before freeze onset. We therefore refer to these neurons as freeze-predictive cells, noting that the predictive nature of their activity is pronounced in Learners compared to Non-Learners and the ANI group.

**Figure 3:**
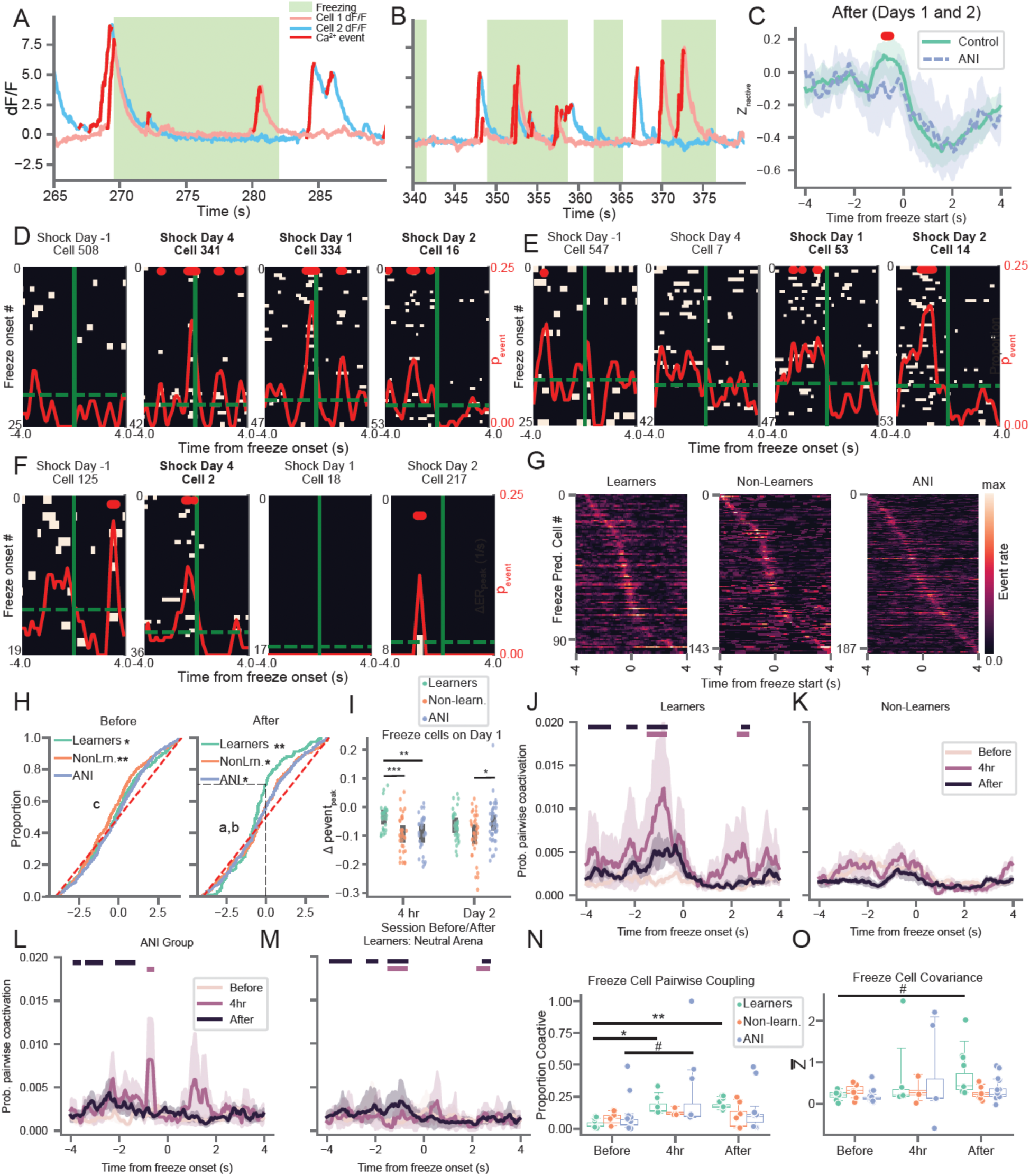
Arresting protein synthesis suppresses the development of coordinated freeze-predicting neural activity. **A) and B)** Example traces from two freeze-predicting cells which exhibit coordinated activity prior to freezing event during the day 1 memory recall session in the shock arena. Red = putative spiking activity, pink = cell shown in C, blue = cell shown in E. **C)** After learning (Days 1 and 2), z-scored population level calcium activity peaks between 0 and 2 seconds prior to freezing for CTRL relative to the ANI group. Line/shading = mean +/- 95% CI. Red: bins with p < 0.05, independent t-test (one-sided, n=7 CTRL mice and 5 ANI mice). **D) and E)** Example Learner freeze-predicting cells identified during the 4 hour (D) or day 1 (E) memory test tracked across sessions. Peri-event calcium activity rasters are centered on freeze onset time (solid green). Dashed green = baseline calcium event probability, red solid = peri-freeze calcium event probability, bins with p < 0.01 (circular permutation test, n=1000) noted with red bars at top. D/E corresponds to pink/blue cells shown in A-B. Bold = session with significant freeze-tuning. **F)** Same as D and E but for ANI mouse freeze-predicting cell identified during the 4 hour session. **G)** Peri-freeze calcium event probability for all freeze-predicting cells detected for each group after learning (Days 1-2), sorted by time of peak activation. **H)** (left) Cumulative distribution of peak peri-freeze activation times before learning. *p=0.49, **p=1e-5 two-sided Wilcoxon rank-sum test, c: p=0.005 Non-Learners v. ANI, 1-sided Mann-Whitney U-test (n = 329/458/543 neurons for Learners/Non-Learners/ANI group) (right) same as left but for after learning. *p < 0.022, **p=2e-7 two-sided Wilcoxon rank-sum test. a: p=0.022 Learners v. Non-Learners, b: p=0.029 Learners v. ANI 1-sided Mann-Whitney U-test (n=194/315/366 neurons for Learners/Non-Learners/ANI group). **I)** Change in peak peri-freeze calcium event probability for all freeze-predicting cells detected during the Day 1 session and either the 4 hour or Day 2 session. p < 0.02 1-way ANOVA each day separately, *p=0.02, **p=0.001, ***p=0.0006 post-hoc Mann-Whiney U-test (n=30/35/29 4h to Day 1 cells and n = 35/37/45 Day 1 to Day 2 cells for Learners/Non-Learners/ANI group. **J)** Pairwise coactivation probability of all freeze-predicting cells for Learners during Before, 4 hour, and After sessions in Shock arena. Maroon/Black bars at top indicate significant increases in coactivation at 4 hour / After time points compared to before, p < 0.05 1-sided Mann-Whitney U-test (n= 4). **K)** Same as J) but for Non-Learners (n=3). **L)** Same as K) but for ANI group (n= 5). **M)** Same as K) but for Learners in Neutral Arena (n=4). **N)** Proportion of freeze-predictive cells with significant pairwise coactivation compared to chance (trial shuffle). *p=4e-8, **p=3.6e-7, #p=0.093. boxplots show population median and 1^st^/3^rd^ quartiles (whiskers, 95% CI) estimated using hierarchical bootstrapping (HB) data with session means overlaid in dots. **O)** Freeze-predicting cells exhibit a trend toward increased peri-freeze covariance (z-scored relative to the Day −2 and −1 covariance values for all cells) for Learners but not Non-Learners or ANI group mice. Mean covariance of freeze-predictive cells from each session shown. #p=0.06. Statistics for K and O: un-paired one-sided HB test after Bonferroni correction, n=10,000 shuffles.

To quantify the reliability of freeze-predictive turning within cells, we tracked the peak, peri-freeze calcium event probability of each freeze cell backwards and forwards in time from the Day 1 recall session, after putative memory consolidation had occurred. We found that freeze-predictive cells exhibited much higher tuning stability spanning from the 4 hour recall to the day 1 recall sessions in Learners than did these cells in other groups (Figure 3I). This indicates that in Learners, reliable freeze-predictive tuning began to emerge between 4 hours and 1 day following learning and was maintained thereafter (see Figure 3E for exemplar cell). In contrast, peri-freezing activities in mice from the ANI group and in Non-Learners were more transient and unreliable during this time span.

Next, we investigated whether the higher reliability of freeze-predicting cells in Learners was related to increased co-activity (see Figure 3A-B) in this neuronal subpopulation by calculating pairwise coactivation of neurons +/- 4 seconds from freeze onset. We found that pairwise coactivation increased significantly from 0-2 seconds prior to freezing during the 4 hour and Day 1-2 sessions for Learners but not for Non-Learners or for ANI group mice (Figure 3J-L). We observed a similar, but much less pronounced, increase for Learners in the Neutral arena for the after learning recall session (Figure 3M). To account for the overall increase in population-level activity prior to freezing (Figure 3C), we calculated chance level pairwise coactivation by shuffling the order of freeze onsets for one cell in each pair 1000 times and determined the probability that the actual pairwise chance exceeded shuffle for each bin. We found that the proportion of freeze-predicting cells with significant coactivation (> 3 consecutive bins with coactivity exceeding 95% of shuffles) increased for Learners, but not Non-Learners or ANI group mice, from before learning to the 4 hour recall session and before learning to after learning recall sessions (Figure 3N).

We then examined whether we observed similar increases in co-activity in the overall neural population. To quantify this, we calculated the covariance between all cell pairs. We found that the covariance of all cell-pairs increased significantly following CFC for Learners for both the 4 hour and after recall sessions (Figure S10H). Furthermore, the observed covariance increase was not driven by freeze-predictive activity (Figure S10J). We observed a similar increase in covariance for the ANI group at the 4hr session and a much smaller, but significant, increase during the after recall sessions (Figure S10H). We observed no such increase for the ANI group when we excluded freezing times from the covariance calculation (Figure S10I). These increases were not observed prior to shock in the neutral arena (Figure S10G), suggesting that the heightened covariance we observed was related to CFC learning. Consistent with our pairwise coactivity analyses, we found a trend toward increase covariance of freeze-predicting cells during the after recall sessions for Learners, but not Non-Learners or ANI mice (Figure 3O). We observed a similar trend in freeze-predicting cell covariance when we downsampled the number of freezing events following learning to match that on days −2 and −1 (Figure S10I).

Finally, we performed additional analyzes focused on the same set of freeze-predictive cells conditioned on their activity during the Day 1 recall test immediately after putative memory consolidation was finalized. Freeze-predicting cells identified on the 1 day recall session displayed a trend toward increased covariance following CFC in Learners but not in Non-Learners or in the ANI group (Figure S10K). However, these cells did not exhibit increased covariance during the previous day’s 4 hour session, suggesting that even though freeze-predictive activity begins to emerge immediately following learning, functional cell connections continue to reorganize up to one day later to form coordinated ensembles. These results demonstrate that freeze-predictive tuning emerges following learning and continues to take shape in the ensuing hours until it stabilizes one day later. Importantly, the coordination of these freeze-predictive cells into neuronal ensembles requires protein synthesis, and they fail to coactivate under memory consolidation failures.

## Discussion

The results of our study provide evidence that HPC spatial and non-spatial representations support contextual memory formation and consolidation. We first probed how protein-synthesis influenced learning-related hippocampal dynamics following CFC. We saw that anisomycin acutely accelerate cell turnover in the 4 hours following learning. In addition to this turnover acceleration, anisomycin’s amnestic effects coincided with a reduction in learning-related remapping, effectively halting HPC contextual representations in their prior state. This paralleled the absence of remapping in the shock arena for Non-Learners, providing strong evidence for the persistence of remapping as a mechanism underlying spatial memory consolidation. Given that remapping *was* observed in Learners during this time window, we speculate that arresting protein synthesis temporarily blocked the formation of new place-fields in the subset of cells that would otherwise undergo remapping following CFC. By this conjecture, existing synapses were weakened to enable the formation of new connections between cells, and these new connections were stifled by anisomycin, diminishing the excitatory drive to these cells to the point where calcium activity was no longer observable. This idea is resonant with the notion that more excitable/active neurons are preferentially involved in memory trace formation (Rashid et al., 2016; Sweis et al., 2021). It also provides an explanation for why we observed accelerated cell turnover following ANI administration as the cells involved in learning are effectively silenced, allowing for a new set of cells to become active. Reduced activity – which may arise from non-specific effects of ANI, or the blockage of constitutive protein translation (Scavuzzo et al., 2019) – could also underlie increased cell turnover. However, our study does not support this view. Consistent with a recent paper which found that ANI did not silence hippocampal neuron activity (Park et al., 2023), we did not observe a widespread shutdown of neural activity as we found that both theta oscillations and sharp-wave ripples were intact in ANI-treated mice.

Surprisingly, remapping occurred in untreated mice that exhibited poor memory recall (Non-Learners); however, this remapping was limited to the neutral arena. This indicates that ANI induced memory failures and poor learning manifest via the same underlying mechanism – a failure to remap in the shock arena – but with important differences. ANI impairs memory by preventing learning-related plasticity which stifles remapping, while poor learning can occur when the shock is improperly associated with the neutral arena, causing remapping therein. In line with previous studies, we found that fear conditioning induced place field remapping in Learners in the shock arena (Wang et al., 2012; Moita et al., 2004). Surprisingly, however, Non-Learners also exhibited remapping, but in the neutral arena only. If remapping is a neural substrate for spatial learning, we speculate that this might explain the increased freezing behavior observed for Non-Learners in the neutral arena: they are improperly associating the neutral arena with shock. Alternatively, the observed remapping could be an indicator of low hippocampal engagement during the task, which has been shown to decrease discrimination between two similar arenas (Wiltgen et al., 2010). However, under either of these situations, we would also expect to observe remapping in the shock arena since Non-Learners freeze equally in both contexts. Future studies could shed further light on this phenomenon. Overall, our findings support the view that the long-term stability of learning-related remapping requires protein synthesis and underlies CFC memory consolidation.

Memory generalization or linking occurs when mice freeze at high levels in a neutral arena. Cai et al. (2016) demonstrated that exposing mice to two arenas five hours apart compared to seven days apart increases the likelihood of memory linking. Despite the close temporal proximity between arena exposures in this experiment, our intent was not to increase contextual fear memory/generalization. Instead, our protocol was designed to produce a moderate level of freezing following conditioning, specifically in the shock arena, as too much freezing would cause insufficient exploration to observe place fields. While close temporal exposure between the conditioning and neutral arena can increase the likelihood of memory linking, it does not guarantee it. Many other factors, such as the amount of pre-exposure time (Frankland et al., 2004) and familiarity with the general experimental procedure can influence contextual memory generalization. It should also be noted that, even procedures designed specifically to create linked/generalized memories produce large variability in freezing levels, with many mice exhibiting low levels of memory linking despite a 5 hour separation between arena exposures (Shen et al., 2022). In contrast, some mice can exhibit generalized freezing in a neutral arena which they never seen before (Wiltgen et al., 2010). The observed variability in memory linking/generalization we observed in our control mice therefore agrees with the high levels of variability observed in previous studies.

Next, we found that a subset of hippocampal cells reliably activated near freezing events during fear memory recall, consistent with previous reports (Schuette et al., 2020; Mocle et al., 2024). These cells activities were particularly heightened 1-2 seconds prior to freezing epochs in animals that exhibited a strong, specific CFC memory but not in the ANI group or untreated mice that exhibited poor memory. The predictive nature of these cells distinguishes them from previously reported immobility cells which occur primarily in CA2 but also in CA1 (Kay et al., 2016). We further found that freeze-predictive cells organized into reliable, co-active neuronal ensembles following learning (Lee & Han, 2022; Rajasethupathy et al., 2015). Importantly, the covariance of all cells increased incrementally for both ANI and Learner groups at the 4 hour recall session, a time point at which protein synthesis is not considered to be required for the induction of (early-phase) long-term potentiation *in vitro* (Frey & Morris, 1997; Nguyen et al., 1994). However, at later time points, when protein synthesis *is* required to maintain (late-phase) long-term potentiation (Frey et al., 1993; Nguyen et al., 1994), between-cell covariance remained high for Learners only, consistent with a recent study demonstrating that the strength of correlated activity in ventral CA1 neurons responsive to a foot shock predicted the strength of contextual fear memory retrieval (Jimenez et al., 2020). It is important to note that all Control mice exhibited heightened calcium activity 1-2 seconds prior to freezing epochs; however, freeze-predictive neurons exhibited higher stability and coactivity for Learners compared to Non-Learners and ANI mice. Therefore, in addition to maintaining learning-related synaptic changes over hours to days, protein synthesis might also be instrumental for producing coordinated firing of large ensembles of neurons to support memory recall.

The synchronous bursts of neuron ensembles we observed prior to freezing may provide a neuronal basis for memory recall. For example, they could indicate the activation of sharp-wave ripples (SWRs): transient, high frequency oscillations observable in the hippocamp local field potential which can facilitate transmitting information throughout the cortex. SWRs are primarily observed during periods of awake immobility and sleep and are hypothesized support memory consolidation and planning through reactivation of neuronal activity patterns observed during learning and active exploration (Buzsáki, 2015). However, SWRs can also occur during periods of movement; these exploratory SWRs (eSWRs) are posited to strengthen connections between place cells for reactivation and subsequent stabilization during sleep (O’Neill et al., 2006). In line with this view, synchronous neuronal activation during eSWRs could strengthen connections between freeze-predicting cells and stabilize their activity patterns to guide future memory recall. Alternatively, coordinated activation of these cells could cooccur with other prominent hippocampal oscillations such as theta or gamma (Colgin, 2016). Whether it occurs with SWRs, theta, or gamma, coincident firing of freeze-predictive ensembles could be important for guiding memory recall. In line with this idea, one study demonstrated that synchronous optogenetic stimulation of engram neurons tagged during learning could artificially reactivate a fear memory even when normal long-term recall of the fear memory was hindered by arresting protein synthesis after learning, suggesting that a memory trace still resided in the network (Ryan et al., 2015). Our findings provide a parsimonious explanation for this previous result by demonstrating that anisomycin halts the co-activation of freeze-predictive cells, weakening the capacity of these neurons to transmit behavior-related information to downstream regions through coincident firing (Lisman, 1997) and impairing their ability to trigger memory recall (Ryan & Frankland, 2022).

Overall, our results provide a bridge between the neuronal activities that underly memory formation and the protein signaling events that are critical for plastic modification of synapses, indicating that protein synthesis is necessary for the formation of new stable spatial representations of an aversive context following learning and for producing coordinated activity of freeze-predictive neurons.

### Limitations of the study

In this study, we casually test how post-learning protein synthesis impacts the neural dynamics related to contextual memory consolidation and recall in the hippocampus. We find that protein synthesis is required for place cells to remap following learning and for freeze-predicting neurons to develop lasting functional connections. We utilized systemic injections of anisomycin to pharmacologically arrest protein synthesis. While anisomycin has been used for decades to block memory consolidation and disrupt long-term potentiation, it produces many off target effects including malaise, which we show in Figure S2, and which could contribute to the accelerated cell turnover and lower cell activity observed in the ANI group following administration. Moreover, it lacks temporal, regional, and cell-type specificity. Therefore, we are unable to determine whether the pharmacologic memory consolidation failure we induce is due to blocking protein synthesis or due to other, non-specific effects, including an acceleration of cell turnover or due to disruptive effects in non-hippocampal brain regions. Future studies could utilize more specific approaches (Shrestha et al., 2020) to disentangle this question. Nonetheless, this study’s finding that anisomycin blocks the coactivity of freeze-predictive cell-pairs is consistent with previous literature demonstrating that protein synthesis is required to produce lasting long-term potentiation between different groups of neurons (Frey and Morris, 1997). We also utilize systemic injections rather than local infusions of anisomycin due to location of a GRIN lens immediately over the hippocampus. The neural and behavioral effects of anisomycin could therefore be due to disruption of protein synthesis in other, non-hippocampal, regions. Regardless, our results are consistent with previous behavioral studies using local hippocampal infusions to block contextual fear memory consolidation and lesion studies demonstrating that the hippocampus is necessary for contextual fear memory formation and consolidation (Kim & Fanselow, 1992; Debiec et al., 2002; Rossatto et al., 2007; Ocampo et al., 2017). Last, we utilize calcium imaging to capture neural activity, which allows for tracking of individual neurons across all stages of the experiment. However, due to the slower temporal dynamics of GCaMP, we are unable to capture fast time-scale hippocampal activity, such as SWRs, reactivation, replay, and theta sequences, that are important for memory formation and consolidation. Future studies could use electrophysiology to explore how these phenomena are likewise impacted by protein synthesis.

## Supporting information

Main and Supplemental Figures

## Acknowledgments

First and foremost, we would like to thank Howard Eichenbaum who helped conceive and design this study before his unfortunate passing in 2017. We would also like to thank Sam McKenzie for his help during early experimental design. We would like to thank Michael Hasselmo and Ian Davison for their support and feedback while performing the recordings for this study. Next, we thank Sam Levy, Dave Sullivan, and Will Mau for their assistance in all phases of calcium imaging throughout. We would like to thank Zach Pennington for valuable feedback concerning anisomycin preparation and administration, and Denise Cai and Lucas Carstensen for analysis suggestions. We would like to thank Pho Hale, Rachel Wahlberg, and Utku Kaya for feedback on the manuscript. We would like to acknowledge the GENIE Program, specifically Vivek Jayaraman, PhD, Douglas S. Kim, PhD, Loren L. Looger, PhD, Karel Svoboda, PhD from the GENIE Project, Janelia Research Campus, Howard Hughes Medical Institute, for providing the GCaMP6f virus. Finally, we would like to acknowledge Inscopix, Inc. for making single-photon calcium imaging miniscopes widely available, and specifically Lara Cardy and Vardhan Dani for all their technical support throughout the experiment. This work was supported by NIH Grants R01 MH052090, R01 MH051570, R01MH117964, NIH NRSA Fellowship 1F32NS117732-01, NIH Early Independence Award DP5 OD023106-01, an NIH Transformative R01 Award, a Young Investigator Grant from the Brain and Behavior Research Foundation, a Ludwig Family Foundation grant, and the McKnight Foundation Memory and Cognitive Disorders award, and Boston University’s Neurophotonics Center.

## Author Contributions

Conceptualization: N.R.K with the help of Howard Eichenbaum; Methodolgy: N.R.K; Software: N.R.K., E.A.R; Validation: N.R.K; Formal Analysis: N.R.K, D.J.O., E.A.R, B.K; Investigation: N.R.K., D.J.O., E.A.R., B.K, S.C.; Data Curation: N.R.K., D.J.O., E.A.R., B.K.; Writing – original draft preparation: N.R.K.; Writing – review and editing; N.R.K, D.J.O, E.A.R., B.K., K.D., S.R.; Visualization: N.R.K.; Project Administration; N.R.K, S.R.; Funding Acquisition: N.R.K., K.D., S.R.

## Materials & Correspondence

All requests for materials and correspondence should be directed to Nat Kinsky or Steve Ramirez.

## STAR★Methods

### Key resources table

| REAGENT or RESOURCE | SOURCE | IDENTIFIER |
| --- | --- | --- |
| Other |  |  |
| pAAV (AAV9).Syn.GCaMP6f.WPRE.SV40 | Addgene | 100837 |
| 1 µL syringe | Hamilton | 65458-02 |
| Micro Injection pump | WPI | UMP3 + Micro4 Controller |
| Microendoscope + Data Acquisition Box | Inscopix | V2 |
| 1028 ch Recording system | Intan | RHS2000 |
| 32 channel headstage | Intan | C3314 |
| 32 channel linear probe | NeuroNexus | A1x32-5mm-50-177 |
| 32 channel uLED probe | Neurolight | Chronic 32ch Optoelectrode |
| Recording cable | Intan | Standard SPI cable |
| Metal microdrive | 3D Neuro | R2Drive |
| Plastic microdrive and rat protective crown | Vöröslakos et al. (2021)<sup>3</sup> | https://doi.org/10.5281/zenodo.8209229 |
| Software |  |  |
| Mosaic 1.2 Image Post-processing software | Inscopix | V1.2 |
| nVista Recording software | Inscopix | nVista |
| OSC1lite Software | This paper | https://doi.org/10.5281/zenodo.8209373 |
| RHX Recording Software | Intan | RHX |
| OpenEphys GUI | OpenEphys | GUI |
| MATLAB | The MathWorks | 2019b |
| TENASPIS: Custom MATLAB code for cell identification and Ca<sup>2+</sup> transient extraction | Mau et al. (2018) | https://github.com/SharpWave/TENASPIS (v4) |
| Custom MATLAB code for cell registration and data analysis | Kinsky et al. (2018) | https://github.com/nkinsky/ImageCamp |
| Custom Python code for Ca<sup>2+</sup> data analysis | This paper | https://github.com/nkinsky/Eraser |
| • Python | Python.org | 3.9.18 |
| • Numpy | Numpy.org | 1.26.3 |
| • Matplotlib | Matplotlib.org | 3.8.2 |
| • Seaborn | Seaborn.pydata.org | 0.13.2 |
| • Pingouin | Pingouin-stats.org | 0.5.4 |
| Custom Python code for electrophysiological data analysis | Kinsky et al. (2024) | https://github.com/diba-lab/NeuroPy |
| • Python | Python.org | 3.11.6 |
| • Numpy | Numpy.org | 1.24.4 |
| • Matplotlib | Matplotlib.org | 3.8.0 |
| • Seaborn | Seaborn.pydata.org | 0.13.0 |
| • Pingouin | Pingouin-stats.org | 0.5.3 |

### Experimental model and study participant details

Sixteen (n = 10 controls, 6 anisomycin) male C57/BL6 mice (Jackson Laboratories), age 16 to 22 weeks during behavioral and imaging experiments and weighing 25-32g were used in this study. Three mice were excluded after performing this study: one mouse after histology revealed the GRIN lens implant and viral expression to be medial to the intended imaging, while the other two were excluded due to unstable/overexpression of GCaMP that produced aberrant calcium activity which emerged toward the end of the experiment. After exclusion of these mice, we retained 8 mice in the control (CTRL) group and 5 mice in the anisomycin group (ANI) group. Additionally, behavioral video tracking files for one CTRL mouse were corrupted during recording during all neutral field recordings from day 0 on: this mouse was excluded from all analyses which required using behavior in the neutral arena (e.g., place field correlations and any analyses where the CTRL group was split into Learners and Non-Learners). Mice were socially housed in a vivarium on a 12 hour light-dark cycle with 1-3 other mice prior to surgery and were housed singly thereafter. Mice were given free access to food and water throughout the study. All procedures were performed in compliance with the guidelines of the Boston University Animal Care and Use Committee.

One male Long Evans rat, 10 months old and weighing 480g, and one female Long Evans rat, 5 months old and weighing 240g, were used for the electrophysiological recordings in this study. Rats were socially housed in a vivarium on an adjusted 12 hour light-dark cycle (lights on at noon, off at midnight) with 1-3 other rats prior to surgery and given free access to food and water throughout the study. One week following recovery from the second surgery (see below), and prior to performing the experiments shown in Figure S5, the second rat was water restricted and performed a separate set of experiments in which she ran for water reward on a linear track during which cells were focally inhibited via delivery of light from one of 12 µLEDs on the implanted silicon probe. During water restriction, her health was monitored and she was weighed daily to ensure she maintained at least 80% of her pre-restriction weight. Following completion of these recordings, the second rat was kept on water restriction while she performed the experiments outlined below in the “Behavioral Paradigm” section under administration of saline or anisomycin. All procedures were performed in compliance with the guidelines of the University of Michigan Animal Care and Use Committee.

### Method details

#### Viral Constructs

For mice experiments we used an AAV9.*Syn*.GcaMP6f.WPRE.SV40 virus from the University of Pennsylvania Vector Core/Addgene with an initial titer of ∼4x10^12^ GC/mL and diluted it into sterilized potassium phosphate buffered saline (KPBS) to a final titer of ∼2-4x10^12^ GC/mL for injection.

For rat experiments, we used an pGP.AAV9.*Syn*.GcaMP7f.WPRE.SV40 virus from the University of Pennsylvania Vector Core/Addgene with an initial titer of 2.6x10^13^ GC/mL and diluted it into sterilized phosphate buffered saline (PBS) to a final titer of 2.6x10^12^ GC/mL for injection. Due to poor expression no imaging was performed.

#### Stereotactic Surgery

We performed two stereotactic surgeries and one base-plate implant on naïve mice, aged 3-8 months, according to previously published procedures (Kinsky et al., 2018; Resendez et al., 2016). Both surgeries were performed under 1-2% isoflurane mixed with oxygen. Mice were given 0.05mL/kg buprenorphine (Buprenex) for analgesia (subcutaneously, SC), 5.0mL/kg of the anti-inflammatory drug Rimadyl (Pfizer, SC), and 400mL/kg of the antibiotic Cefazolin (Pfizer, SC) immediately after induction. They were carefully monitored to ensure they never dropped below 80% of their pre-operative weight during convalescence and received the same dosage of Buprenex, Cefazolin, and Rimadyl twice daily for three days following surgery. In the first surgery, a small craniotomy was performed at AP −2.0, ML +1.5 (right) and 250nL of GcaMP6f virus (at the titer noted below) was injected 1.5mm below the brain surface at 40nL/min using a 1µL Hamilton syringe and infusion pump. The needle remained in place a minimum of 10 minutes after the infusion finished at which point it was slowly removed, the mouse’s scalp was sutured, and the mouse was removed from anesthesia and allowed to recover.

3-4 weeks after viral infusion, mice underwent second surgery to attach a gradient index (GRIN) lens (GRINtech, 1mm x 4mm). After performing an ∼2mm craniotomy around the implant area, we carefully aspirated cortex using blunted 25ga and 27ga needles under constant irrigation with cold, sterile saline until we visually identified the medial-lateral striations of the corpus callosum. We carefully removed these striations using a blunted 31ga needle while leaving the underlying anterior-posterior striations intact, after which we applied gelfoam for 5-10 minutes to stop any bleeding. We then lowered the GRIN lens to 1.1mm below bregma. Note that this entailed pushing down ∼50-300µm to counteract brain swelling during surgery. We then applied Kwik-Sil (World Precision Instruments) to provide a seal between skull and GRIN lens and then cemented the GRIN lens in place with Metabond (Parkell), covered it in a layer of Kwik-Cast (World Precision Instruments), and then removed the animal from anesthesia and allowed him to recover after removing any sharp edges remaining from dried Metabond with a dental drill and providing any necessary sutures.

Finally, after ∼2-4 weeks we performed a procedure in which the mouse was put under light anesthesia to attach a base plate for easy future attachment of a miniature epifluorescence microscope (Ghosh et al., 2011, Inscopix, Inc.). Importantly, no tissue was cut during this procedure. After induction, we attached the base plate to the camera via a set screw, set the camera’s focus level at ∼1/3 from the bottom of its range, and carefully lowered the camera objective and aligned it to the GRIN lens by eye, and visualized fluorescence via nVistaHD until we observed clear vasculature and putative cell bodies expressing GcaMP6f (Resendez et al., 2016). To counteract downward shrinking during curing, we then raised the camera up ∼50µm before applying Flow-It ALC Flowable Composite (Pentron) between the underside of the baseplate and the cured Metabond on the mouse’s skull. After light curing, we applied opaque Metabond over the Flow-It ALC epoxy to the sides of the baseplate to provide additional strength and to block ambient light infiltration. Mice were allowed to recover for several days prior to habituation to camera attachment and performance of the behavioral task outlined below. In the event that we did not observe clear vasculature and cell bodies when we first visualized fluorescence we covered the GRIN lens with Kwik-Cast and removed the mouse from anesthesia without attaching the baseplate. We then waited an additional week and repeated the steps above.

For rats, we performed two surgeries in a similar manner as described above for mice. However, rats were administered pre-operative and post-operative Meloxicam orally for analgesia (in lieu of Buprenex) and triple-antibiotic was applied locally (in lieu of Cefazolin injections) to the incision at the end of surgery. Meloxicam was additionally administered for two days post-surgery during recovery, and animals were monitored daily for a minimum of seven days during recovery. 0.4mL of a lidocaine/bupivacaine cocktail were given under the scalp to provide local anesthesia at the incision site. In the first surgery, 1000nL of GCaMP7f virus was infused in the prelimbic cortex at the center of a 1mm craniotomy (AP + 2.9, ML + 3.6, from Bregma, DV −3.0 at an 18 degree angle from top of brain). Following infusion, ∼1.5 mm of overlying cortex was removed and a 23ga needle was lowered to ∼500µm above the target site. Then, a 0.6 x 7 mm GRIN lens was lowered to 3.0mm below the top of the brain, the area between the skull and lens was sealed with Kwik-Sil, and the lens was affixed to the skull with Vivid-Flow light-curable composite (Pearson Dental) and Metabond (Parkell). The lens was then covered in Kwik-Sil for protection. During this surgery, ground and reference screws were also placed over the cerebellum and a 3d printed crown base was attached to the rat’s skull (Vöröslakos et al., 2021) to which crown walls and top were connected and to further protect the lens and future microdrive/probe implant. The rat was screened for fluorescence 8-12 weeks later, but no cell dynamics were observed so no imaging equipment was implanted for this rat.

16 weeks later, the rat was again given pre-operative Meloxicam and anesthetized under isoflurane for probe implant (Kinsky et al., 2023). The crown walls were removed and a 1.0mm craniotomy was performed at AP-4.8, ML+3.6 from bregma. After removing dura and stopping bleeding with cold, sterile saline, a NeuroNexus A1x32-5mm-50-177 probe, attached to a metal microdrive, was implanted at 2.3 mm below the brain surface and the metal drive base was attached to the skull with Unifast light cured dental epoxy (Henry-Schein). The craniotomy was sealed with Dow-Sil, the probe was protected with dental wax, and the ground and reference wires were connected to the probe electronic interface board (EIB). The crown walls were re-attached, the EIB was connected to the crown walls, and the rat was removed from isoflurane and allowed to recovery. The rat was monitored daily for 7 days prior to recording, during which the probe was lowered ∼1mm until sharp wave ripples and spiking activity were visualized indicating localization of the probe in the CA1 cell layer. Full details including videos demonstrating the implant process are also documented in Kinsky et al., (2023).

#### Histology procedures

Hippocampal slices were prepared following extraction from mice in accordance with the standard methods and guidelines of the Boston University Animal Care and Use Committee. In brief, mice were euthanized with Euthasol (Virbac), transcardially perfused with paraformaldehyde (PFA), and decapitated. Following extraction, brains were placed in PFA for approximately 48 hours before undergoing sectioning. Brains that were sliced using a Cryostat underwent an additional step of sucrose cryoprotection and subsequent freezing in -80C. Brains were mounted to the slicing platform using Tissue Tek O. C. T. (Sakura) and kept at −30C throughout sectioning. 50µm slices were collected across the entire aspiration site in the dorsal hippocampus region. Brains that were sliced using a vibratome were stabilized using super glue and submerged in 1% PBS. A Leica VT1000 S vibratome was equipped with a platinum coated double edged blade (Electron Microscopy Sciences, Cat. #72003-01) and set to a maximal speed of 0.9mm/s for collecting 50 µm slices. Slices prepared from both the cryostat and microtome were directly mounted onto (type of slides go here) and cover-slipped using DAPI following sectioning. No histology was performed in the rat study.

For cell death (apoptosis) experiments, adult male mice weighing 20.1-25g were injected with either anisomycin (150mg/kg, IP, ∼0.11mL) or saline (IP, ∼0.11mL) and perfused 4hr later with PBS followed by 4% paraformaldehyde. Brain tissue was removed and fixed for 48hr in PFA. Fixed tissue was sliced into 50μm thick free-floating sections and stored into 0.01% azide. Tissue was sliced coronally (50um thickness) on a vibratome and stained using Biosensis (Ready-to-Dilute) Fluoro-Jade C (FJC) staining kit. Hippocampal sections were selected and mounted on charged slides and left to dry overnight for proper adhesion. Following kit instructions, slides incubated in a coplin jar with 9 parts of 80% ethanol and 1 part of sodium hydroxide for 5 minutes. Slides were then rinsed in 70% ethanol for 2 minutes followed by 2 minutes in distilled water. To minimize background fluorescence, slides were submerged in 9 part distilled water and 1 part potassium permanganate for 5 minutes, followed by 2 x 2 minute rinses in distilled water. Slides were then incubated in 9 parts distilled water and 1 part FJC in the dark for 15 minutes, followed by 3 x 1 minute rinses in distilled water. Finally, tissue dried for 10-20 minutes before being submerged in xylene for 10 minutes and mounted with DPX. Fluorescence was visualized using a confocal microscope.

#### Behavioral Paradigm

Prior to surgery mice were handled to habituate them subsequent camera attachment. 3-7 days following base plate attachment surgery we conditioned mice to the imaging procedures by further handling them for 5-10 minutes for a minimum three days. During this handling a plastic “dummy” microscope (Inscopix) of approximately the same size/weight as the imaging camera was attached to each mouse’s head and remained on his head for 1-2 hours in his home cage. When it became easy to attach the scope to the mouse’s head a real imaging miniscope was attached to head and an optimal focus plane chosen. We then recorded three 5 minute imaging videos at this focus and +/- ¼ turn (∼25μm) in the mouse’s home cage. These movies were processed as described in the Image Acquisition and Processing section and an optimal zoom was chosen based on whichever focus plane maximized cell yield and produced clear looking cell bodies. Animals were then placed in a novel environment with a different size and shape compared to the experimental environments for a 10 minute session to habituation them to the general experimental outline and ensure that they explored novel arenas.

Following habituation to the imaging procedures mice performed a contextual fear conditioning (CFC) task with simultaneous imaging of hippocampus neurons over the course of 10 days. Note that all recording sessions are referred to by their time relative to applying the mild foot-shock and the arena in which the recording occurred: e.g., Shock Day −2 occurred in the shock arena two days prior to foot-shock while Neutral 4 hours occurred four hours after foot-shock in the neutral arena. A typical day (Days −2, −1, 1, 2, and 7) consisted of two separate 10 minute recording blocks/sessions: one in the Neural arena and one in the shock arena. Mice first explored a square (neutral) arena, placed in the center of a well-lit room, for 10 minutes. The neutral arena was a square constructed of 3/8” plywood (25cm x 25cm x 15 cm), which was painted yellow with sealable paint. Additionally, one wall was painted with black horizontal stripes for visual orientation purposes. The neutral arena was wiped down with 70% ethanol ∼10 minutes prior to recording. After 10 minutes of exploration the experimenter took the mouse out of the arena, leaving the miniscope camera on their head and placed the mouse in its home cage on a moveable cart upon which it was immediately transported down a short hallway to second room. Therefore, the time between the end of the neutral and shock arena recordings was approximately five minutes.

The second room was dimly lit and contained the fear conditioning (shock) arena. The shock arena (Coulbourn Instruments, Whitehall, PA, USA) consisted of metal-panel side walls, Plexiglas front and rear walls, and a stainless-steel grid floor composed of 16 grid bars (22cm x 22cm). Following 10 minutes of exploration of the shock arena, mice were removed from the arena, the camera was removed, and mice were returned to their home cage. Both arenas were wiped down with 70% ethanol ∼10min prior to recording to eliminate any odor cues. Note that mice always explored the neutral arena first and the shock arena second. For the Day 0 sessions, mice first explored the neutral arena for 10 minutes and were transported to the shock arena as usual. However, during this session (Shock Day 0) the mouse was immediately given a single 0.25mA shock and allowed to explore the arena for an additional 60 seconds only before being removed and returned to his cage. Efficacy of shock was confirmed post-hoc by eye by the presence of jumping/darting behavior immediately post-shock. The 4 hour session was identical to the Day −2, 1, 1, 2, and 7 sessions. With the exception of the 4 hour session, all recording sessions were performed in the first half of the mouse’s life cycle while the 4 hour session occurred in the second half of the light cycle.

On day zero, after the camera was removed and prior to returning to their home cage, mice received an intraperitoneal injection of either anisomycin (150 mg/kg, Sigma-Aldrich A9789) or the equivalent amount of vehicle. After injection, they were returned to their cage for 4 hours until the next recording session began.

Following extensive habituation to a rest box during the seven day recovery period, rat neural activity and behavior was recorded across ∼ 5 hours. Following a 15-30 minute baseline recording (PRE) in the rest box, the animal was given an I.P. injection of anisomycin and then immediately placed on a novel linear track which he explored for 45 minutes (TRACK). The rat was then placed back into the rest box for 3.5 hours (POST). Following that, the animal was placed on a second novel track for 45 minutes (TRACK2) followed by a brief recording in the rest box (POST2). The second rat underwent a similar procedure, except that there was a 2 hour POST recording prior to the TRACK recording and no POST2 recording. Similar procedures were followed the day before and after anisomycin injection but using saline for injection instead of anisomycin. Data was acquired continuously throughout with the exceptions of periodic cable disconnections to perform the I.P. injection, start a new recording epoch, and disconnect/reconnect cables that became twisted.

#### Anisomycin

For mice recordings, 25 mg of anisomycin (Sigma Aldrich) was dissolved into 50 µL of 6N HCl and 500 µL of 1.8%NaCl. ∼125 µL of 1N NaOH was then added to the solution followed by 0.1-0.5 µL of 1N NaOH, testing pH after each addition until a final pH of 7.0 to 7.5 was reached, with a final concentration of 24-27 mg/mL. In the case that pH rose above 7.5 during titration and and/or the anisomycin went back into precipitate, small amounts (10-20 µL) of 6N HCl were added until particles were no longer visible and the titration with 1N NaOH was restarted. Mice were administered 150mg/kg of anisomycin solution via intraperitoneal injection, or ∼0.15-0.18mL for a typical 30g mouse.

For rat recordings, 100mg of anisomycin was dissolved into 1.6mL of 0.1N HCl (in 0.9% saline). ∼240 µL of 1N HCl was added, then 10-12 µL of NaOH was added in 1-2 µL amounts, testing pH between each step until a pH of 7-7.5 was reached. 0.9% Saline was then added until the appropriate concentration was reached to inject 1.5mL of anisomycin at 150mg/kg. In one rat (Rat1 in Figure S5), due to a small amount of waste, the final amount injected corresponded to 100 mg/kg.

### Quantification and statistical analysis

#### Behavioral Tracking and Fear Metrics

We utilized two different camera/software configurations for tracking animal behavior. Both configurations generated a TTL pulse at the beginning of behavioral tracking to synchronize with image acquisition. We utilized Cineplex software (v2, Plexon) to track animal location at 30Hz in the neutral arena. We used FreezeFrame (Actimetrics) to track animal location in the shock arena at 3.75Hz. Animal location was obtained post-hoc via custom-written, freely available Python software (www.github.com/wmau/FearReinstatement). We observed inconsistent frame rates and inaccurate acquisition of behavioral video frames for one mouse in the neutral arena during the day 0, 4 hour, and day 1-2 sessions. These sessions were excluded from analysis.

Freezing was calculated by first downsampling neutral position data to 3.75 Hz to match the sample rate used in the shock arena. We then identified freezing epochs as any periods of 10 consecutive frames (2.67 seconds) or more where the mouse’s velocity was less than 1.5cm/second.

#### Neural Discrimination

We evaluated the extent to which each animal’s behavior reflected the expression of a context-specific fear memory through a behavioral discrimination index (DI_beh_), calculated as follows:

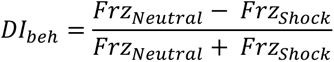

Where Frz_Neutral_ and Frz_Shock_ are the percentages of time spent freezing in the neutral and shock arenas, respectively. Thus, a negative DI_beh_ value indicated more freezing behavior in the shock arena (indicating successful encoding of a context-specific fear memory), a positive DI_beh_ value indicated more freezing behavior in the neutral arena, and a DI_beh_ value around zero indicated equal/low freezing behavior in each arena (indicating the formation of a non-specific or weak fear memory).

#### Imaging Acquisition and Processing

Brain imaging data was obtained using nVista HD (Inscopix) at 720 x 540 pixels and a 20 Hz sample rate. Note that imaging data for one mouse was obtained at 10 Hz. Prior to neuron/calcium event identification we first pre-processed each movie using Inscopix Imaging Suite (Inscopix) software. Preprocessing entailed three steps a) motion corrections, and b) cropping the motion-corrected movie to eliminate any dead pixels or areas with no calcium activity, and c) extracting a minimum projection of the pre-processed movie for later neuron registration. We did not analyze one imaging session in which we had to reconnect the camera cable mid-session and could not synchronize the imaging data with behavioral data. Maximum projections of imaging movies were made using the Inscopix Imaging Suite or custom-written functions based off of an open-source MATLAB library (Muir & Kampa, 2015).

#### Electrophysiological Recordings

Data was acquired using an Intan 1028 channel recording system through OpenEphys software into binary format and behavior was tracked using Optitrack high resolution cameras and Motive image acquisition software.

#### Data Analysis

Data analysis was performed in both Python and MATLAB software. Python analysis code is available at https://github.com/nkinsky/Eraser.

#### Spike sorting and analysis

Electrophysiological recordings were automatically clustered using SpyKING CIRCUS software (Yger et al., 2018) and units were manually curated in phy (https://github.com/cortex-lab/phy/). Units were grouped into single units if they exhibited a clear refractory period and were well-isolated from other putative spikes. Other units which exhibited a clear waveform but were either poorly isolated or exhibited refractory period violations were classified as multi-unit activity (MUA). All single units and MUA were combined and cross-correlograms for the combined activity were created for each epoch of the recording separately.

#### Tenaspis

Neuron regions-of-interest (ROIs) and calcium events were identified using a custom written, open source algorithm employed in MATLAB 2016b called A Technique for Extracting Neuronal Activity from Single Photon Neuronal Image Sequences (Tenaspis) (Mau et al., 2018). This procedure was comprehensively documented in Kinsky et al., 2018: “Tenaspis is open-source and available at: https://github.com/SharpWave/TENASPIS. First, Tenaspis filters each calcium imaging movie with a band-pass filter per (Kitamura et al., 2015) to accentuate the separation between overlapping calcium events. Specifically, Tenaspis smooths the movie with a 4.5 μm disk filter and divides it by another movie smoothed with a 23.6 μm disk filter. Second, it adaptively thresholds each imaging frame to identify separable pockets of calcium activity, designated as blobs, on each frame. Blobs of activity are accepted at this stage of processing only if they approximate the size and shape of a mouse hippocampal neuron, as measured by their radius (min = ∼6μm, max = ∼11μm), the ratio of long to short axes (max = 2), and solidity (min = 0.95), a metric used by the *regionprops* function of MATLAB we employ to exclude jagged/strange shaped blobs. Third, Tenaspis strings together blobs on successive frames to identify potential calcium transients and their spatial activity patterns. Fourth, Tenaspis searches for any transients that could results from staggered activity of two neighboring neurons. It rejects any transients whose centroid travels more than 2.5μm between frames and whose duration is less than 0.20 seconds. Fifth, Tenaspis identifies the probable spatial origin of each transient by constructing putative regions-of-interest (ROIs), defined as all connected pixels that are active on at least 50% of the frames in the transient. Sixth, Tenaspis creates initial neuron ROIs by merging putative transient ROIs that are discontinuous in time but occur in the same location. Specifically, it first attempts to merge all ROIs whose centroids are less than a distance threshold of ∼0.6μm from each other. In order to merge two transient ROIs, the two-dimensional Spearman correlation between the ROIs must yield r^2^ > 0.2 and p < 0.01. Tenaspis then successively increases the distance threshold and again attempts to merge ROIs until no more valid merges occur (at a distance threshold of ∼3μm, typically). Seventh, Tenaspis integrates the fluorescence value of each neuron ROI identified in the previous step across all frames to get that neuron’s calcium trace, and then identifies putative spiking epochs for each neuron. Specifically, it first identifies the rising epochs of any transients identified in earlier steps. Then, it attempts to identify any missed transients as regions of the calcium trace that have a) a minimum peak amplitude > 1/3 of the transients identified in step 3, b) a high correlation (p < 0.00001) between active pixels and the pixels of the average neuron ROI identified in step 6, and b) a positive slope lasting at least 0.2 seconds. Last, Tenaspis searches for any neuron ROIs that overlap more than 50% and whose calcium traces are similar and merges their traces and ROIs.”

#### Between Session Neuron Registration

We utilized custom-written, freely available MATLAB code (available at https://github.com/nkinsky/ImageCamp) to perform neuron registration across sessions in accordance with previous results (Kinsky et al., 2018). The details of this procedure described in Kinsky et al. (2018) are reproduced here:

“Neuron registration occurred in two steps: session registration and neuron registration.

##### Session registration

Prior to mapping neurons between sessions, we determined how much the imaging window shifted between sessions. In order to isolate consistent features of the imaging plane for each mouse (such as vasculature or coagulated blood), we created a minimum projection of all of the frames of the motion-corrected and cropped brain imaging movie for each recording session. One session (“registered session”) was then registered to a base session using the “imregtform” function from the MATLAB Image Processing Toolbox, assuming a rigid geometric transform (rotation and translation only) between images, and the calculated transformation object was saved for future use.

##### Neuron Registration

Next, each ROI in the registered session was transformed to its corresponding location in the base session. Each neuron in the base session was then mapped to the neuron with the closest center-of-mass in the registered session, unless the closest neuron exceeded our maximum distance threshold of 3 pixels (3.3 μm). In this case the base session neuron was designated to map to no other neurons in the registered session. If, due to high density of neurons in a given area, we found that multiple neurons from the base session mapped to the same neuron in the registered session, we then calculated the spatial correlation (Spearman) between each pair of ROIs and designated the base session ROI with the highest correlation as mapping to the registered session ROI.

For multiple session registrations, the same procedure as above was performed for each session in two different ways. First, we registered each session directly to the first session in the experiment and updated ROI locations/added new ROIs to set of existing ROIs with each registration. This helped account for slight day-to-day drift in neurons ROIs due to shifts in vasculature, build-up of fluid underneath the viewing window, creep/shrinkage of dental cement, etc. Second, to ensure that neuron ROIs did not drift excessively across sessions we also performed all the above steps but did NOT update ROI locations allowing us to register each set of ROIs to those furthest away chronologically. The resulting mappings were then compared across all sessions, and any neuron mappings that differed between the two methods (e.g., ROIs that moved excessively across the duration of the experiment) were excluded from analysis. Those that remained in the same location were included.”

The procedure to assess the quality of across session registration was described in Kinsky et al. (2018) and is reproduced here: “We checked the quality of neuron registration between each session-pair in two ways: 1) by plotting the distribution of changes in ROI orientations between session and comparing it to chance, calculated by shuffling neuron identity between session 1000 times, and 2) plotting ROIs of all neurons between two sessions and looking for systematic shifts in neuron ROIs that could lead to false negatives/positives in the registration.” All session-pairs (except those few in which we could not synchronized imaging and behavioral data as noted above) met the above two criteria and were thus included in our analysis.

Cells that had calcium activity in the first session (neutral arena) for which we did not identify a matching neuron in the second session (shock arena) were classified as OFF cells. Likewise, neurons active in the shock arena with no matching partner in the neutral arena were classified as ON cells.

All neuron registrations were cross validated by overlaying ROIs from each session and evaluating their match by eye. In a few cases, we noticed erroneous registrations and adjusted our between-session neuron alignment by calculating the rigid geometric transformation using 4-5 cell ROIs active in both sessions. Figure S11 provides examples of our between-session neuron registration results and quantification procedures.

#### Neural Discrimination Metrics

The extent to which gross hippocampal ensemble activity differed between arenas was calculated in two ways. First, we calculated the proportion of cells that turned ON and OFF between arenas divided by the total number of cells active in either arena.

Next, we calculated the extent to which each neuron active in both arenas distinguished between arenas by changing its event rate in a manner analogous to DI_beh._ However, we took the absolute value to account for the fact that both positive and negative event rate changes could reflect neural differentiation between arenas. Then, we took the mean across all neurons to obtain a neural discrimination index (DI_Neuron_):

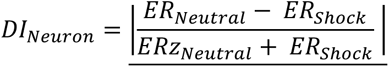

#### Generalized Linear Model (GLM) to Assess anisomycin effects on the number of active cells

All GLM analyses were performed using the *GLM* function in the Python *statsmodels* package. The dependent variable was the number of neurons recorded in each session, normalized to the amount recorded on Day −2. Covariates included were a constant/intercept term, group (ANI vs. CTRL), arena (Shock vs. Neutral), anisomycin stage (acute = 4 hour, after = Days 1, 2, and 7), freezing ratio, and two interaction terms: group x acute and group x after. The GLM was fit assuming a gaussian distribution with the identify link function and using the iteratively reweighted least squares method.

#### Placefield Analysis

Calcium transients occurring when the mouse was running greater than or equal to 1.5cm/second were spatially binned (1cm by 1cm) and occupancy normalized following which place fields were identified and quantified in a manner similar to Kinsky et al. (2018), reproduced here:

“Spatial mutual information (SI) was computed from the following equations, adapted from (Olypher et al., 2003)

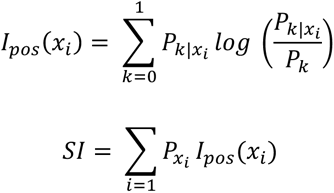

where:

_-_ P_xi_ is the probability the mouse is in pixel *x_i_*
- P_k_ is the probability of observing *k* calcium events (0 or 1)
_-_ P_k|xi_ is the conditional probability of observing *k* calcium events in pixel *x_i_*.

The SI was then calculated 1000 times using shuffled calcium event timestamps, and a neuron was classified as a place cell if it 1) had at least 5 calcium transients during the session, and 2) the neuron’s SI exceeded 95% of the shuffled Sis…We defined the extent of a place field as all connected occupancy bins whose smoothed event rate exceeded 50% of the peak event rate occupancy bin.”

Since spatial mutual information is biased by the number of samples (Olypher et al., 2003), we re-sampled the behavioral tracking data to match that of the imaging data (20Hz). This required up-sampling the shock arena data (3.75Hz->20Hz) and down sampling the neutral arena data (30Hz->20Hz).

Placefield similarity between sessions was assessed by first smoothing the 2-d occupancy normalized event rate maps with a gaussian kernel (2.5cm std), flattening the smoothed maps into a vector, and then performing a Spearman correlation between all neurons active in both sessions. To quantify chance-level place field similarity we randomly shuffled the mapping between neurons from the first to the second session before performing the Spearman correlation. We then repeated this procedure 100 times.

To assess the possibility that the configuration of place fields rotated together coherently between sessions (Kinsky et al., 2018), we again performed a Spearman correlation but after rotating the 2-d occupancy map in the second session 90 degrees. Since, due to small camera distortions, some 2-d occupancy maps were not square, one some occasions we resized (minimally) the second map to match the size/shape of the first map using the reshape function in Python’s *numpy* package prior to correlate the two maps. We repeated this in successive 90 degree increments and then took the mean correlation of all neurons that were active in both sessions to determine the optimal/“best” rotation of the place field map as that which maximized the correlation between sessions.

We also performed a “center-out” rotation analysis to assess coherent place field rotations between sessions. First, the angle to the pixel with the maximum occupancy normalized event rate was identified for each cell. Second, this angle was recalculated for the same cell in a different session in the same box. These two angles were subtracted to get the “center-out” rotation between sessions. Sessions which exhibited a coherent rotation displayed a peak in a histogram of center-out angles at 0, 90, 180, or 270 degrees, while sessions which exhibited global remapping exhibited a uniform distribution of rotation angles.

#### Freeze-predicting Cell Analysis

Freeze onset and offset times were first identified for each mouse/session as noted in the *Behavioral Tracking and Fear Metrics* section above. We then formed calcium event rasters using the neural activity for each cell +/- 4 seconds from freeze onset, organizing the data into a *nfreeze_onsets x ntime_bins* array. We then summed this raster along the 0^th^ dimension to get a freeze tuning curve. To calculated significance, we randomly, circularly shifted the putative spiking activity for a cell and calculated a shuffled tuning curve in a similar manner to the actual data. We repeated this procedure 1000 times, and calculated significance for each time bin as the number of shuffles where the shuffled tuning exceeded the actual tuning curve divided by 1000. Last, we designated cells as significantly freeze-tuned if they had 3 or more bins with p < 0.01 and were active on at least 25% of freezing events.

#### Covariance Analysis

Putative spiking activity for each cell was first binned into 0.5 second windows and z-scored after binning, forming a *ncells x nbins* array. The covariance of this array was then calculated using the *cov* function in numpy, returning a *ncells x ncells* array. For between-session comparisons, cells active in both sessions were registered and a new array was formed with the base (1^st^) session covariance in the lower diagonal and the registered (2^nd^) session in the upper diagonal. All entries along the main diagonal were ignored. This analysis was also performed including only peri-freeze times (to assess peri-freeze covariance), after randomly downsampling the number of freeze epochs to match the average from days −2 and −1 (to control for increased sampling on days 1 and 2), and after excluding all peri-freeze times (to assess non peri-freeze covariance).

#### Coactivation Analysis

Putative spiking activity was first binned relative to the start of each freezing epoch (“freeze onset”), yielding a *nfreeze_onsets x nbins* array of binarized neural activity (1 = active, 0 = inactive). Pairwise coactivation was then calculated by taking the dot product of the binarized neural activity arrays between each pair of cells and taking the mean along the first axis, yielding the probability that both cells in a given cell-pair were active in each sampling bin. To calculate population-level coactivity, the number of active cells was first calculated for each recording session. The average number of active cells was then calculated across all peri-freeze trials (+/- 4 seconds) and this number was z-scored relative to activity across the entire session. To determine which cell-pairs exhibited significant pairwise coactivity beyond chance level, we first formed a *nfreeze_onsets x nbins* array of neural activity for each cell in a pair. We then randomly permuted the rows of the array for the second cell in the pair and calculated a shuffled level coactivation probability for that pair as described above. We repeated this process 100 times for each cell pair, and then calculated a p-value for each bin as 1 – (# of bins where shuffle exceeds actual coactivity) / 100. A cell-pair was designated to have significant pairwise coactivation if it had > 3 consecutive bins with p < 0.05.

#### Hierarchical Bootstrapping

We utilized hierarchical bootstrapping (HB, Saravanan et al., 2020) to estimate confidence intervals and P-values for different metrics in line with code found at https://github.com/soberlab/Hierarchical-Bootstrap-Paper. For each metric we generated a population of 10,000 values by resampling with replacement at each level of the data hierarchy (first mouse, then for each mouse, one session, then the variable measured, e.g. place field correlations or freeze-predicting cell covariance). We then pooled all the values and calculated the test statistic (typically the mean) and corresponding confidence interval / interquartile range. One or two-tailed tests were used with ⍺ = 0.05, and the p-value was calculated by using the joint probability distribution of the bootstrapped samples to determine that the mean of one group was greater than the mean of the other group. All hierarchical bootstrapped data was visualized using boxplots (mean and 1^st^/3^rd^ quartiles) with whiskers extending to the 95% confidence intervals created using the boxplot function from the Python package *Seaborn*. Chance level was calculated by shuffling cell identity before calculating each metric, and the mean and 95% confidence intervals were visualized using the *matplotlib* package.

#### Parametric statistics

Where applicable, all parametric statistics used are noted in the corresponding figure legend.

## FIGURE LEGENDS

**Figure S1: CTRL animals exhibit behavioral variability following learning and similar rates of cell turnover between arenas vs. across days.**

**A)** CTRL mice freezing on all days. Red = shock arena, blue = neutral arena. **B)** Distribution of DI_beh_ scores for all mice in CTRL group on days 1 and 2. Dashed line indicates cutoff between Learners and Non-Learners. **C)** Cell overlap 1 day apart in the same arena (days −2 to −1 and 1 to 2) vs. cell overlap between arenas on the same day (days −2, −1, 1, 2) for all mice. *p=4.05e-7, ρ=0.81 Spearman correlation, n=26 overlap values across 13 mice. **D)** Same as C) but for neural discrimination index. *p=6.29e-7, ρ=0.82 Spearman correlation. **E)** Freeze ratio plots with each mouse’s value connected by a line. **F)** No difference in thigmotaxis prior to conditioning in Neutral arena, dots show mean thigmotaxis ratio for each mouse from Days −2 and −1, p=0.23 ANOVA. **G)** Same as F) but in Shock arena, p=0.95 ANOVA. **H)** Neutral arena freezing ratio for each session after shock plotted versus Neutral arena freezing on day of training (day 0) with Pearson correlation and associated p-value (two-sided) shown, n = 7 mice. **I)** Same as H) but for Shock arena freezing versus Neutral arena freezing on day of training.

**Figure S2: Non-specific effects of Anisomycin include a reduction in locomotion**

**A)** 4 mice were given I.P injections of anisomycin only (no shock) and their locomotion was tracked over 24 hours. Normal activity did not return to baseline until between 6 and 24 hours later. Black solid/dashed lines = 4 hour mean +/- std freezing ratio for non-ANI fear conditioned mice shock arena (see B). **B)** Freezing ratios for all mice in the Shock arena prior to conditioning and 4 hours after conditioning shown for reference. **C)** Freezing ratios in Neutral arena immediately before and one day after conditioning. Lines show mean +/- std.

**Figure S3: Absolute cell numbers recorded across all sessions**.

**A)** Total number of neurons recorded across all sessions in Control group. **B)** Same as A but for ANI group. **C)** Same as A but for Learners. **D)** Same as A but for Non-Learners. **E)** Histogram of cell counts across all sessions with each group mean shown with dashed lines.

**Figure S4: Anisomycin does not cause neuronal cell death at 4hr post injection in the hippocampus.**

**A)** Experiment schematic. Animals were given I.P. injections of either ANI (n=8) or saline (n=7). 4 hours later, brains were extracted and processed for Fluoro-Jade C. **B)** Fluoro-Jade C positive neuronal count normalized to area; **C-H)** Representative images of whole hippocampus (HPC), CA1, and dentate gyrus (DG) in **C-E)** anisomycin-injected mice or **F-H)** saline-injected mice. Scale bar equals 200μm.

**Figure S5: Reduced activity following anisomycin administration is not an imaging artifact and does not result from global disruption of electrical neural activity in hippocampal neurons.**

**A-C)** Neural activity was tracked across ∼5 hours before and after systemic administration of anisomycin in a rat. **A)** Cross correlograms for all single and multi-unit activity combined are shown from the pre epoch in a rest box (15 minutes), running on a novel track immediately following anisomycin injection (45 minutes), post epoch in the rest box (3.5 hours), running on a second novel track (45 minutes), and a second post epoch in the rest box (15 minutes). Clear modulation of firing at the theta timescale is observed. **B)** Example trace from electrode in pyramidal cell layer of CA1 showing theta activity 10 minutes and 4 hours 15 minutes post injection anisomycin injection. **C)** Example sharp wave ripple events occurring from 25 minutes to 4 hours 15 minutes post anisomycin injection across 9 channels of a linear probe spanning above to below the pyramidal cell layer. **D-F)** Same as A-C but for a different rat following systemic anisomycin injection. F) Shows one trace from an electrode in the cell layer with a raster of spiking activity from all units recorded shown below the trace, demonstrating a population burst coincident with each sharp wave ripple. **G-I**) Same as D-F but the following day after systemic saline (control) injection demonstrating no lasting effects of anisomycin 24 hours after injection. **J-K)** The signal-to-noise ratio of all mouse neurons captured using calcium imaging and active between sessions was tracked between sessions. **J)** Mean height of calcium transient peaks for all cells matched from day −1 to 4 hour session. p>0.63 both groups, two-sided t-test (n=8 CTRL and 7 ANI, 1 additional CTRL mouse included whose video tracking behavioral data was corrupted and could therefore could not be classified as a Learner or Non-Learner). **K)** Same as J) but tracking cells from day −1 to day 1, p>0.68 both groups, two-sided t-test.

**Figure S6**: **Coherent Place Field Rotations Observed Between Sessions**

**A)** Example animal trajectories from neutral arena day −2 (top row) and day −1 (middle row) with calcium activity overlaid (red). Each column corresponds to one cell. Bottom row shows data rotated 90 degrees, demonstrating a coherent rotation of spatial activity for all neurons. **B)** Smoothed, occupancy normalized calcium event maps corresponding to data shown in A). **C)** The angle from the center of the arena to each cell’s maximum intensity place field center was calculated for each session (center-out angle). The distribution of center-out angles plotted, demonstrating a coherent rotation of place fields from Day −2 to Day −1 by 90 degrees. **D)** Place field correlations (smoothed event maps) between sessions indicate apparently low stability across days without considering rotations, giving the false impression that the place field map randomly reorganizes between sessions. **E)** High correlations were observed after considering a coherent 90 degree rotation between sessions, indicating that place fields retain the same relative structure but rotate together as a whole. **F)** Mean correlations for each mouse without considering rotations gives the impression of instability before/after shock and heightened remapping for all groups from before to after learning. Boxplots show population median and 1^st^/3^rd^ quartiles (whiskers, 95% CI) estimated using hierarchical bootstrapping (HB) data with session means overlaid in dots. Dashed line and grey shading show mean and 95% CI of correlations calculated from shuffling cell identify 1000 times between sessions.

**Figure S7: Example stable and remapping place fields across sessions**

**A)** Example place field plots for Learner mouse in Neutral arena Before conditioning (Day −2 to Day −1), from Before to After conditioning (Day −1 to Day 1), and After consolidation (Day 1 to Day 2) demonstrating high stability in Neutral arena across all time points. **(top row)** Mouse trajectory in black with cell calcium activity overlaid in red. **(bottom row)** Smoothed, occupancy normalized calcium event spatial maps corresponding to raw trajectory and event data shown in top row, with warmer colors indicating high event rates and cool colors indicating low event rates. Spearman correlation value between event rate maps shown at top. Dashed lines denote separate different cells at each time point, and solid lines separate different comparison times. **B)** Same as A) but for different Learner in Shock arena demonstrating remapping. Red box shows two example remapping cells in the Shock arena. **C)** and **D)** Same and A) and B) but for two different Non-Learners with red box showing remapping cells in the Neutral Arena. **E)** and **F)** Same as A) and B) but for two different ANI group mice showing stable place fields between sessions at all time points.

**Figure S8: Place field correlations with STM (4 hour) session**

**A)** Place field correlations for all mice combined for Day −1 vs 4 hour session. Boxplots show population median and 1^st^/3^rd^ quartiles (whiskers, 95% CI) estimated using hierarchical bootstrapping (HB) data with session means overlaid in dots. Dashed line and grey shading show mean and 95% CI of correlations calculated from shuffling cell identity 1000 times between sessions. **B)** Same as A) but for 4 hour vs Day 1 session. All hierarchical bootstrap test comparisons are ns. **C)** Place field correlations for all mice in each group from before to after shock (Day −1 to Day 1). Boxplots show population median and 1^st^/3^rd^ quartiles (whiskers, 95% CI) estimated using hierarchical bootstrapping (HB) data with session means overlaid in dots. Significant p-values calculated using a one-sided HB permutation test are shown directly on each panel.

**Figure S9: Population Vector (PV) correlations.**

**A)-E)** 1D PV correlations between sessions including only cells active in BOTH sessions. **A)** Before (Day −2 vs −1), *p=0.042, ***p<0.0006. **B)** After (Day 1 vs 2) *p=0.034, #p=0.072. **C)** Before v After (Day −1 vs 1), *p<0.039, **p=0.0098. **D)** Day −1 vs 4 hr session. **E)** 4 hr session vs Day 1 *p=0.0049, #p=0.076. **F)** Day 2 vs Day 7, *p=0.0189, **p=0.0164, #p=0.064. Green = Learners, Orange = Non-Learners, Blue = ANI. Boxplots show population median and 1^st^/3^rd^ quartiles (whiskers, 95% CI) estimated using hierarchical bootstrapping (HB) data with session means overlaid in dots. Dashed line and grey shading show mean and 95% CI of correlations calculated from shuffling cell identify 1000 times between sessions. Statistics: un-paired one-sided HB test after Bonferroni correction.

**Figure S10: Freeze-cell covariance increases are driven by Learners and not purely a result of less freezing in Non-Learners and the ANI group.**

**A)-C)** Example freeze-predicting cells tracked across sessions forward and backward in time from the day indicated in bold. Peri-event calcium activity rasters are centered on freeze onset time (solid green). Dashed green = baseline calcium event probability, red solid = peri-freeze calcium event probability, bins with p<0.01 (circular permutation test) noted with red bars at top. D/E corresponds to pink/blue cells shown in A-B. **A)** Example freeze-predictive cell from Non-Learner **B)-C)** Example freeze-predicting cells from two different Learners. **D)** Proportion freeze-predicting cells detected in Neutral arena across days. Bars=mean, line=std. **E)** Same as D) but for Shock arena. **F)** After learning (Days 1 and 2), z-scored population level calcium activity peaks between 0 and 2 seconds prior to freezing for both Learners and Non-Learners. Line/shading = mean +/- 95% CI. Red: bins with p < 0.05, independent t-test (one-sided, n=4 Learners and 3 Non-Learners). **G)** Mean covariance of all cells in neutral arena prior to learning exhibit small changes, compare y-axis to Figure 3O and S10H, H.**p=0.0048, ***p<1e-8, #p=0.06. **H)** Same as G) but for all cells in shock arena. *p=0.026, **p=0.00052, ***p<2.5e-6. **I)** Same as G) but for freeze predictive cells only, peri-freeze times only, and after randomly downsampling the number of freeze events to match the average number observed during days −2 and −1. #p=0.056 **J)** Same as H) but excluding all peri-freeze times. *p=0.0018**p=1.2e-4, ***p=2.1e-5. **K)** Mean covariance of freeze-predicting cells detected on Day 1 tracked across time. #p=0.068. For G-K, boxplots show population median and 1^st^/3^rd^ quartiles (whiskers, 95% CI) estimated using hierarchical bootstrapping (HB) data with session means overlaid in dots. Dashed line and grey shading show mean and 95% CI of correlations calculated from shuffling cell identify 1000 times between sessions. Statistics: un-paired one-sided HB test after Bonferroni correction. **L-M)** There are no significant differences between the proportion of active units (z-scored) peaks for Control animals compared to the ANI group between 0 and 2 seconds prior to freezing before conditioning (L) and at the 4 hour session (M). Line/shading = mean +/- 95% CI. Red: bins with p < 0.05, independent t-test (two-sided).

**Figure S11: Between Session Neuron Registration Metrics**

**A)** Example neuron registration between the Day −1 and 4 Hour session for one mouse in the shock arena depicting cell ROIs active during the Day −1 session only (blue), the 4 Hour session only (teal), and both the Day −1 and 4 Hour session (yellow) with ROIs which were successfully registered across sessions outlined in red. Insets show magnified ROIs and demonstrate that cells registered across days exhibit similar shape and orientation. **B)** and **C)** Minimum projections of the imaging movie from the sessions shown in A demonstrating high day-to-day stability, evidenced by clear alignment of landmarks (e.g. vasculature) between sessions. **D)** Change in ROI orientation for all neurons registered to the Day −2 session for the mouse shown in A). Note that the majority of registered neurons exhibit very small changes in orientation between sessions, even up to 9 days later (Day −2 to Day 7). **E)** and **F)** Similar to **D)** but for all mice in the CTRL and ANI groups separately for Day −1 to Day 4 (from before to after anisomycin administration). Note a similar distribution of ROI orientation changes, indicating that the observed acceleration of cell turnover following anisomycin administration is not due to poor neuron registration. **G)** Side-by-side comparison of all neuron ROI changes for each group shown as a cumulative density function.

