## Supplementary material for "Erasable Hippocampal Neural Signatures Predict Memory Discrimination": Main and Supplemental Figures

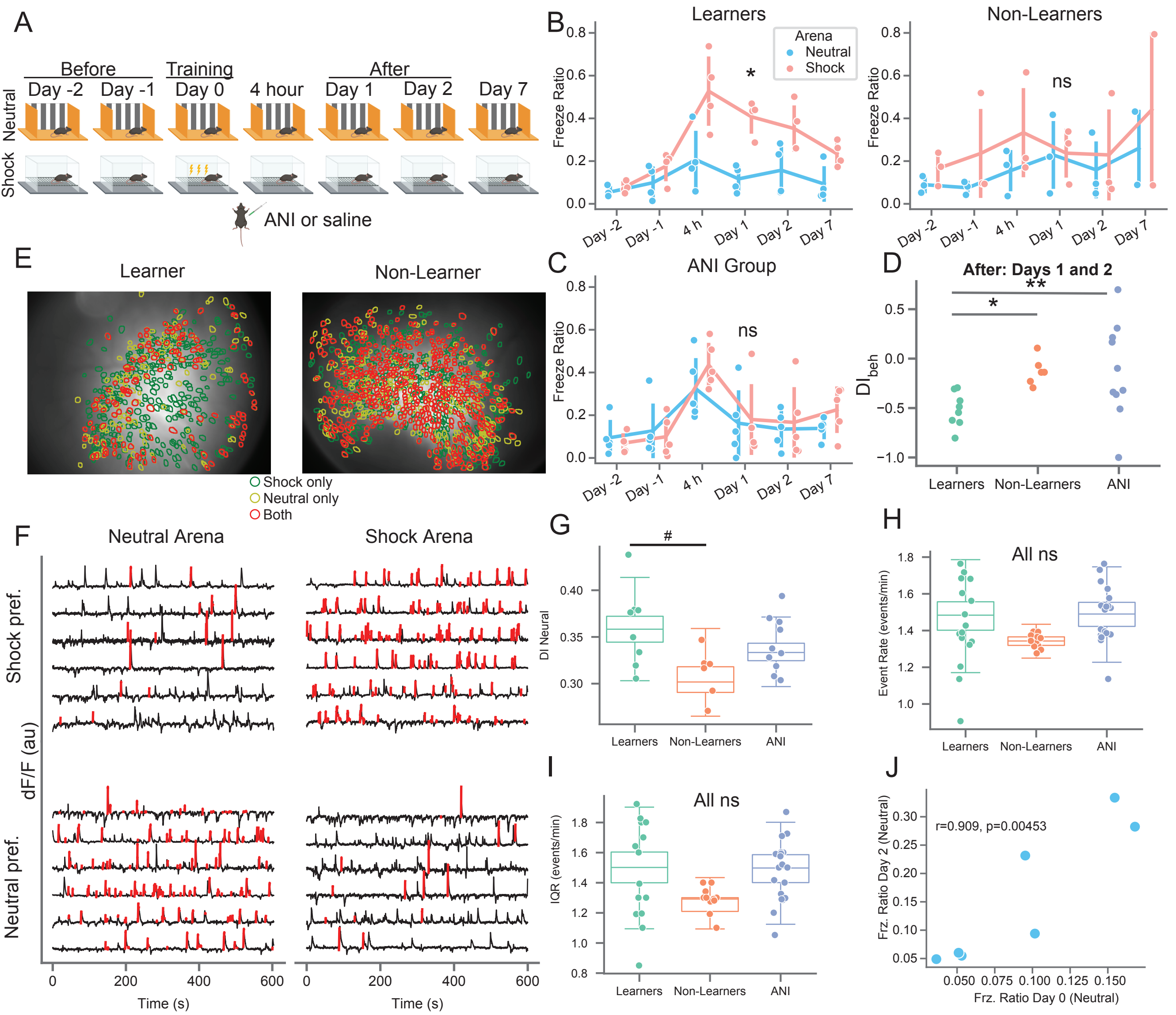


**Figure 1: Mice exhibit variability in memory recall and neural activity prior to learning in a contextual fear conditioning task.** **A)** Schematic of the behavioral paradigm**.** Mice freely explored two distinct arenas (neutral and shock) for 10 minutes each day. Mice underwent mild contextual fear conditioning on day 0 in the shock arena followed by immediate I.P. administration of anisomycin or vehicle in their home cage. Memory recall tests were conducted 4 hours and 1, 2, and 7 days post-shock. The time of each session is referenced to the shock session. **B)** (left) Learner (CTRL) mice freezing on all days. Red = shock arena, blue = neutral arena. *p=4.5e-0.4 shock **–** neutral freezing from day-1 to day 1 one-sided paired t-test (n=4 mice, t=13.4). (right) Same but for Non-Learner (CTRL) mice (n=3 mice, p=0.249, t=0.819). **C)** Same as B but for ANI group (n=5 mice, p=0.219, t=0.859). **D)** Behavioral discrimination between arenas after shock (Days 1-2) shows formation of a specific fear memory for Learners only, by definition (positive = more freezing in neutral arena, negative = more freezing in shock arena, 0 = equal freezing in both arenas). *p=0.009 (t=3.56), #p=0.06 (t=1.83) 1-sided t-test of mean DI value from Days 1 & 2, n=8/6/10 sessions for Learners/Non-Learners/ANI group. **E)** (left) Neural overlap plots between neutral and shock arenas for an example Learner mouse on day -1, before shock. Green = cells active in the shock arena only, yellow = cells active in the neutral arena only, orange = cells active in both arenas. (right) Same for example Non-Learner on day -2 showing higher overlap of active cells between arenas. **F)** Example calcium activity from the Learner mouse shown in C (left) for cells active in both arenas. Black = calcium trace, red = putative spiking activity during transient rises. Top row shows shock arena preferring cells, bottom row shows neutral arena preferring cells. **G)** Neural discrimination index (DIneural) between groups on Days -2 and -1. Boxplots show population median and 1st/3rd quartiles (whiskers, 95% CI) estimated using hierarchical bootstrapping (HB) data with session means overlaid in dots, #p=0.09 after Bonferroni correction for multiple comparisons. **H)** Same as G) but for event rate in Shock arena. **I)** Same as G) but for event rate interquartile range (IQR) in Shock arena. **J)** Freezing in Neutral arena on Day 2 vs. Neutral arena on Day 0. Pearson correlation value and p-value (two-sided) shown on plot. Statistics for G-J: un-paired one-sided HB test for days -2 and -1 after Bonferroni correction, n=10,000 bootstraps.

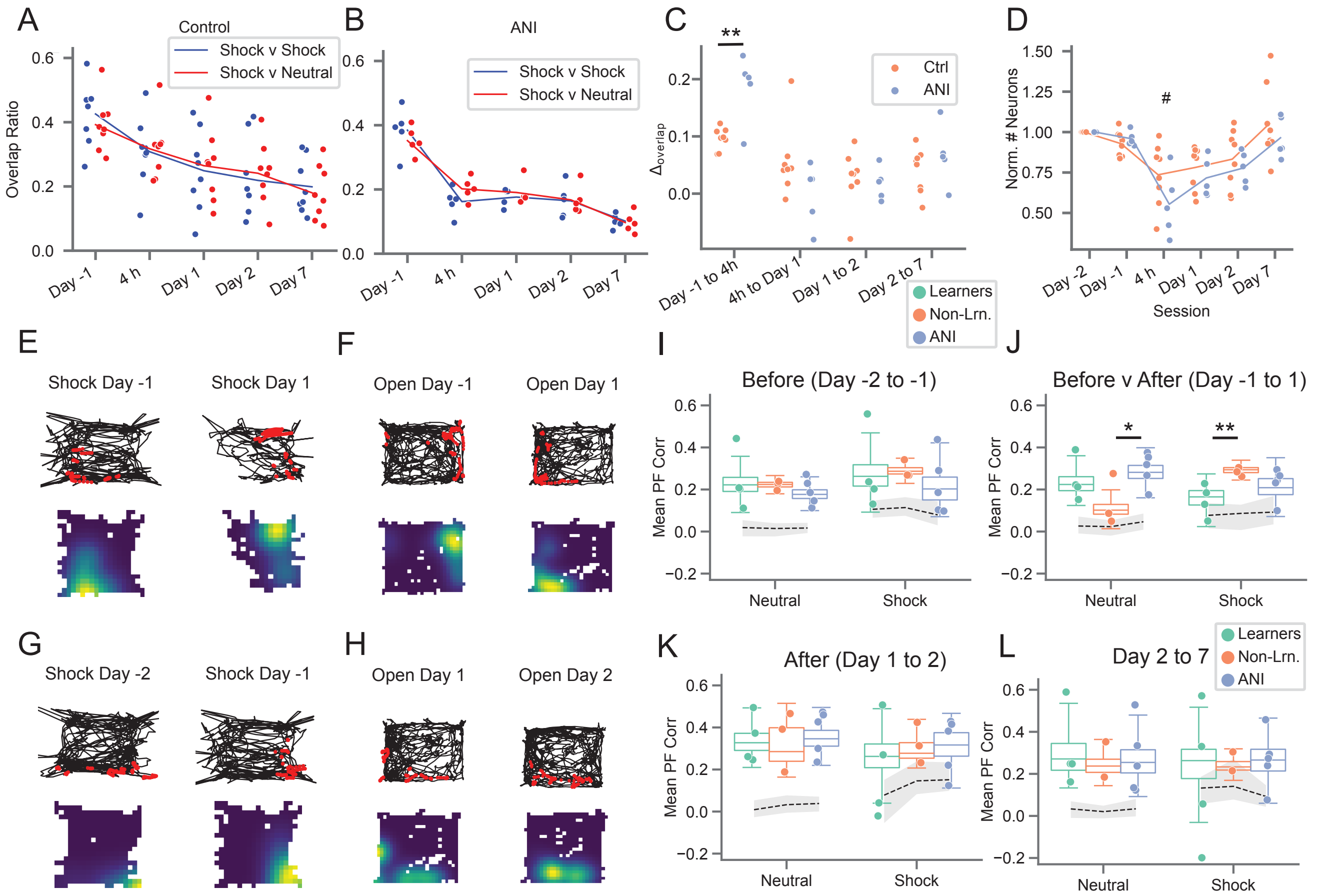


**Figure 2: Preventing protein synthesis accelerates cell turnover and stifles learning-related place field remapping. A)** Cell overlap ratio with Day -2 session, CTRL group. Blue = within shock arena, red = shock v. neutral arena. **B)** Same as A) but for ANI group. **C)** Change in overlap ratios from A) and B), dots show values from both arenas for each mouse, **p=0.00174 two-sided t-test of mean value for each mouse (t=4.11, n=7 CTRL mice and 5 ANI mice). **D)** Number of active neurons observed each day, normalized to day -1. p=2.e-5 freeze-ratio, #p=0.056 group x 4 hr session interaction, p=0.094 group x after interaction, generalized linear model. **E)** and **F)** Example place fields exhibiting learning-related remapping. **E)** Place field in shock arena from Learner mouse. (top) Example mouse trajectory (black) with calcium activity (red) overlaid for the same cell from day -1 to 1 in shock arena, (bottom) occupancy normalized rate maps for the same cells with warm colors indicating areas of high calcium activity. **F)** Same as E) but for Non-Learner mouse in Neutral arena. **G)** and **H)** Example stable place fields. **G)** Same as E) but for a different cell from same mouse in the shock arena prior to conditioning. **H)** Same as F) but for a different cell from the same mouse in the neutral arena after conditioning. **I)** Place field correlations for all mice before shock (Days -2 and -1), boxplots show population median and 1st/3rd quartiles (whiskers, 95% CI) estimated using hierarchical bootstrapping (HB) data with session means overlaid in dots. Dashed line and grey shading show mean and 95% CI of correlations calculated from shuffling cell identify 1000 times between sessions. **J)** Same as I) but for Day -1 to Day 1, *p=0.0496, **p=0.0034. **K)** Same as I) but for Day 1 to Day 2. **L)** Same as I) but for Day 2 to Day 7. Statistics for I-L: un-paired one-sided HB test after Bonferroni correction, n=10,000 bootstraps.

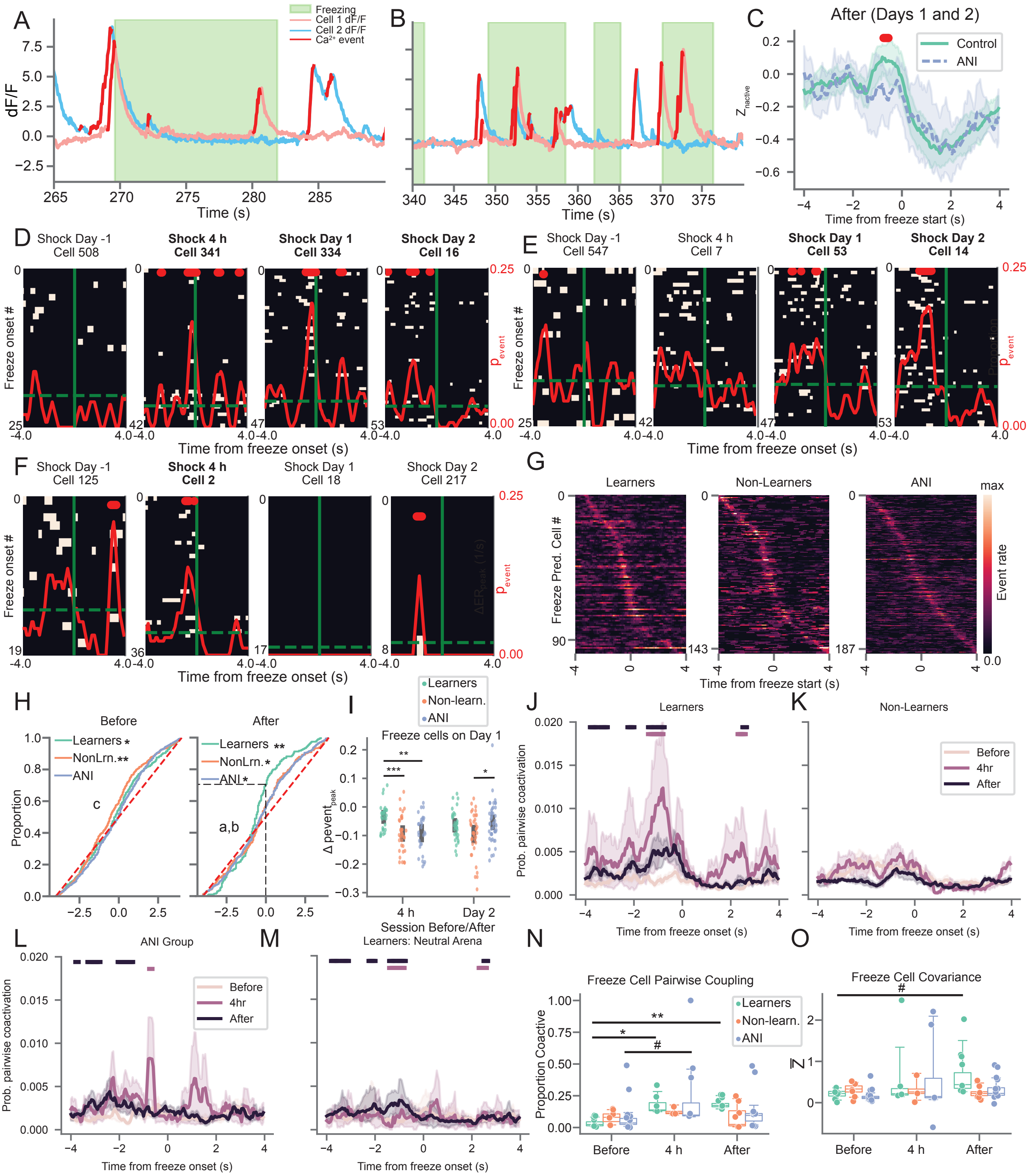


**Figure 3: Arresting protein synthesis suppresses the development of coordinated freeze-predicting neural activity. A)** and **B)** Example traces from two freeze-predicting cells which exhibit coordinated activity prior to freezing event during the day 1 memory recall session in the shock arena. Red = putative spiking activity, pink = cell shown in C, blue = cell shown in E. **C)** After learning (Days 1 and 2), z-scored population level calcium activity peaks between 0 and 2 seconds prior to freezing for CTRL relative to the ANI group. Line/shading = mean +/- 95% CI. Red: bins with $p < 0.05$, independent t-test (one-sided, n=7 CTRL mice and 5 ANI mice). **D)** and **E)** Example Learner freeze-predicting cells identified during the 4 hour (D) or day 1 (E) memory test tracked across sessions. Peri-event calcium activity rasters are centered on freeze onset time (solid green). Dashed green = baseline calcium event probability, red solid = peri-freeze calcium event probability, bins with $p<0.01$ (circular permutation test, n=1000) noted with red bars at top. D/E corresponds to pink/blue cells shown in A-B. Bold = session with significant freeze-tuning. **F)** Same as D and E but for ANI mouse freeze-predicting cell identified during the 4 hour session. **G)** Peri-freeze calcium event probability for all freeze-predicting cells detected for each group after learning (Days 1-2), sorted by time of peak activation. **H)** (left) Cumulative distribution of peak peri-freeze activation times before learning. *p=0.49, **p=1e-5 two-sided Wilcoxon rank-sum test , c: p=0.005 Non-Learners v. ANI, 1-sided Mann-Whitney U-test (n = 329/458/543 neurons for Learners/Non-Learners/ANI group) (right) same as left but for after learning. *$p < 0.022$, **p=2e-7 two-sided Wilcoxon rank-sum test. a: p=0.022 Learners v. Non-Learners, b: p=0.029 Learners v. ANI 1-sided Mann-Whitney U-test (n=194/315/366 neurons for Learners/Non-Learners/ANI group). **I)** Change in peak peri-freeze calcium event probability for all freeze-predicting cells detected during the Day 1 session and either the 4 hour or Day 2 session. $p < 0.02$ 1-way ANOVA each day separately, *p=0.02, **p=0.001, ***p=0.0006 post-hoc Mann-Whiney U-test (n=30/35/29 4h to Day 1 cells and n = 35/37/45 Day 1 to Day 2 cells for Learners/Non-Learners/ANI group. **J)** Pairwise coactivation probability of all freeze-predicting cells for Learners during Before, 4 hour, and After sessions in Shock arena. Maroon/Black bars at top indicate significant increases in coactivation at 4 hour / After time points compared to before, $p< 0.05$ 1-sided Mann-Whitney U-test (n= 4). **K)** Same as J) but for Non-Learners (n=3). **L)** Same as K) but for ANI group (n= 5). **M)** Same as K) but for Learners in Neutral Arena (n=4). **N)** Proportion of freeze-predictive cells with significant pairwise coactivation compared to chance (trial shuffle). *p=4e-8, **p=3.6e-7, #p=0.093. boxplots show population median and 1st/3rd quartiles (whiskers, 95% CI) estimated using hierarchical bootstrapping (HB) data with session means overlaid in dots. **O)** Freeze-predicting cells exhibit a trend toward increased peri-freeze covariance (z-scored relative to the Day -2 and -1 covariance values for all cells) for Learners but not Non-Learners or ANI group mice. Mean covariance of freeze-predictive cells from each session shown. #p=0.06. Statistics for K and O: un-paired one-sided HB test after Bonferroni correction, n=10,000 shuffles.

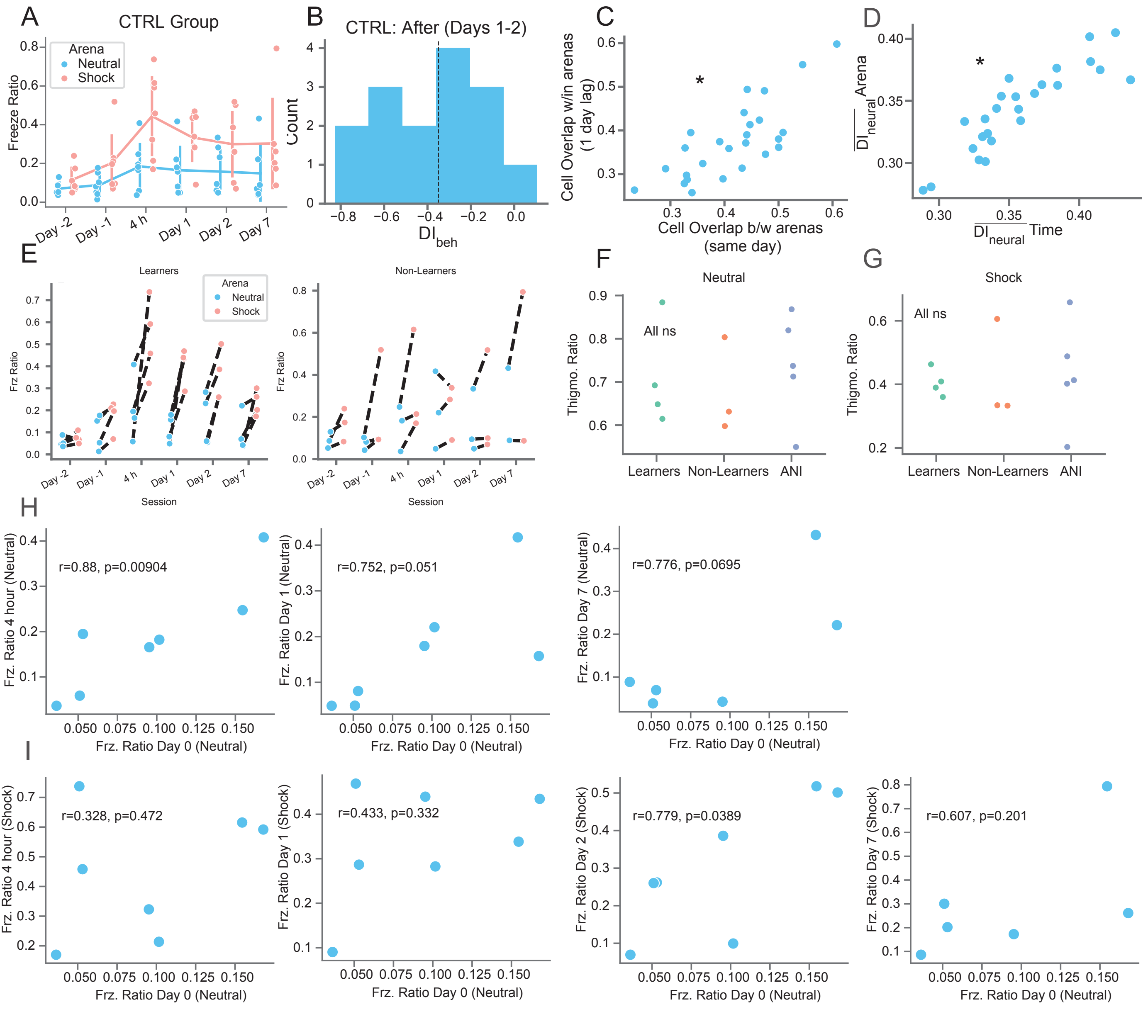


**Figure S1: CTRL animals exhibit behavioral variability following learning and similar rates of cell turnover between arenas vs. across days.**

**A)** CTRL mice freezing on all days. Red = shock arena, blue = neutral arena. **B)** Distribution of DI¨beh scores for all mice in CTRL group on days 1 and 2. Dashed line indicates cutoff between Learners and Non-Learners. **C)** Cell overlap 1 day apart in the same arena (days -2 to -1 and 1 to 2) vs. cell overlap between arenas on the same day (days -2, -1, 1, 2) for all mice in CTRL group. *p=1.7e-5, =0.74 Spearman correlation. **D)** Same as C) but for neural discrimination index. *p=2.35e-8, =0.56 Spearman correlation. **E)** Freeze ratio plots with each mouse's value connected by a line. **F)** No difference in thigmotaxis prior to conditioning in Neutral arena, dots show mean thigmotaxis ratio for each mouse from Days -2 and -1, p=0.23 ANOVA. **G)** Same as F) but in Shock arena, p=0.95 ANOVA. **H)** Neutral arena freezing ratio for each session after shock plotted versus Neutral arena freezing on day of training (day 0) with Pearson correlation and associated p-value (two-sided) shown. **I)** Same as H) but for Shock arena freezing versus Neutral arena freezing on day of training.

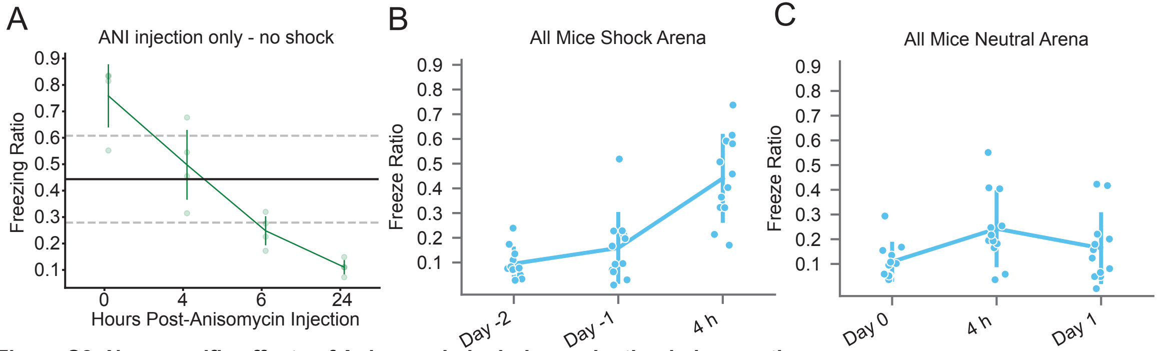


**Figure S2: Non-specific effects of Anisomycin include a reduction in locomotion**
**A)** 4 mice were given I.P injections of anisomycin only (no shock) and their locomotion was tracked over 24 hours. Normal activity did not return to baseline until between 6 and 24 hours later. Black solid/dashed lines = 4 hour mean +/- std freezing ratio for non-ANI fear conditioned mice shock arena (see B). **B)** Freezing ratios for all mice in the Shock arena prior to conditioning and 4 hours after conditioning shown for reference. **C)** Freezing ratios in Neutral arena immediately before and one day after conditioning. Lines show mean +/- std.

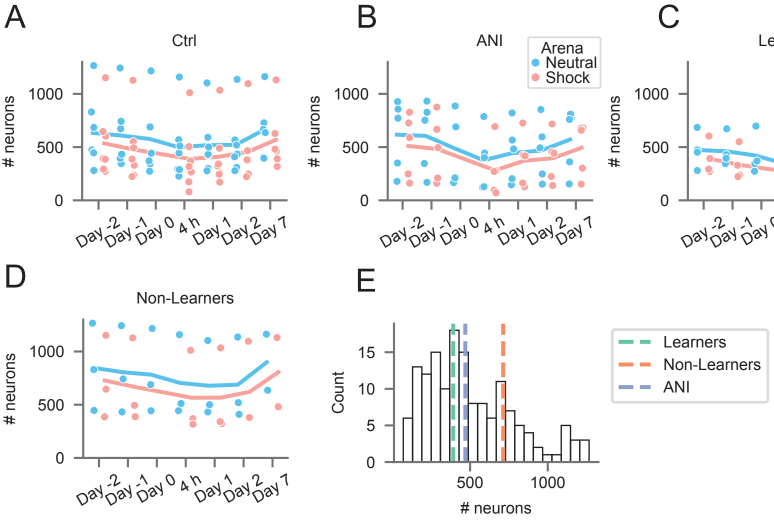


**Figure S3: Absolute cell numbers recorded across all sessions.**
**A)** Total number of neurons recorded across all sessions in Control group. **B)** Same as A but for ANI group. **C)** Same as A but for Learners. **D)** Same as A but for Non-Learners. **E)** Histogram of cell counts across all sessions with each group mean shown with dashed lines.

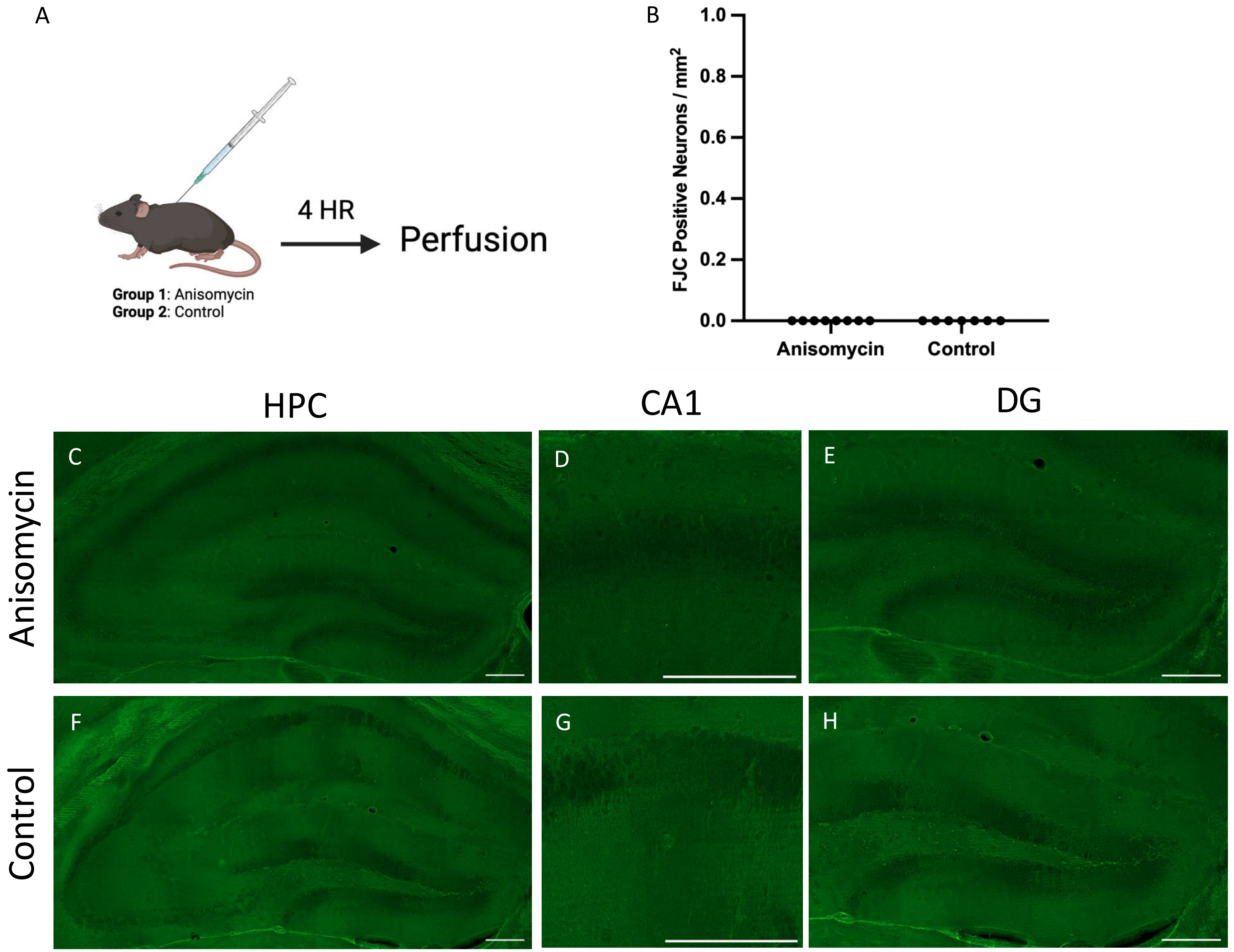


**Figure S4: Anisomycin does not cause neuronal cell death at 4hr post injection in the hippocampus.**
**A)** Experiment schematic. Animals were given I.P. injections of either ANI (n=8) or saline (n=7). 4 hours later, brains were extracted and processed for Fluoro-Jade C. **B)** Fluoro-Jade C positive neuronal count normalized to area. **C-H)** Representative images of whole hippocampus (HPC), CA1, and dentate gyrus (DG) in **C-E)** anisomycin-injected mice or **F-H)** saline- injected mice. Scale bar equals 200Ķm.

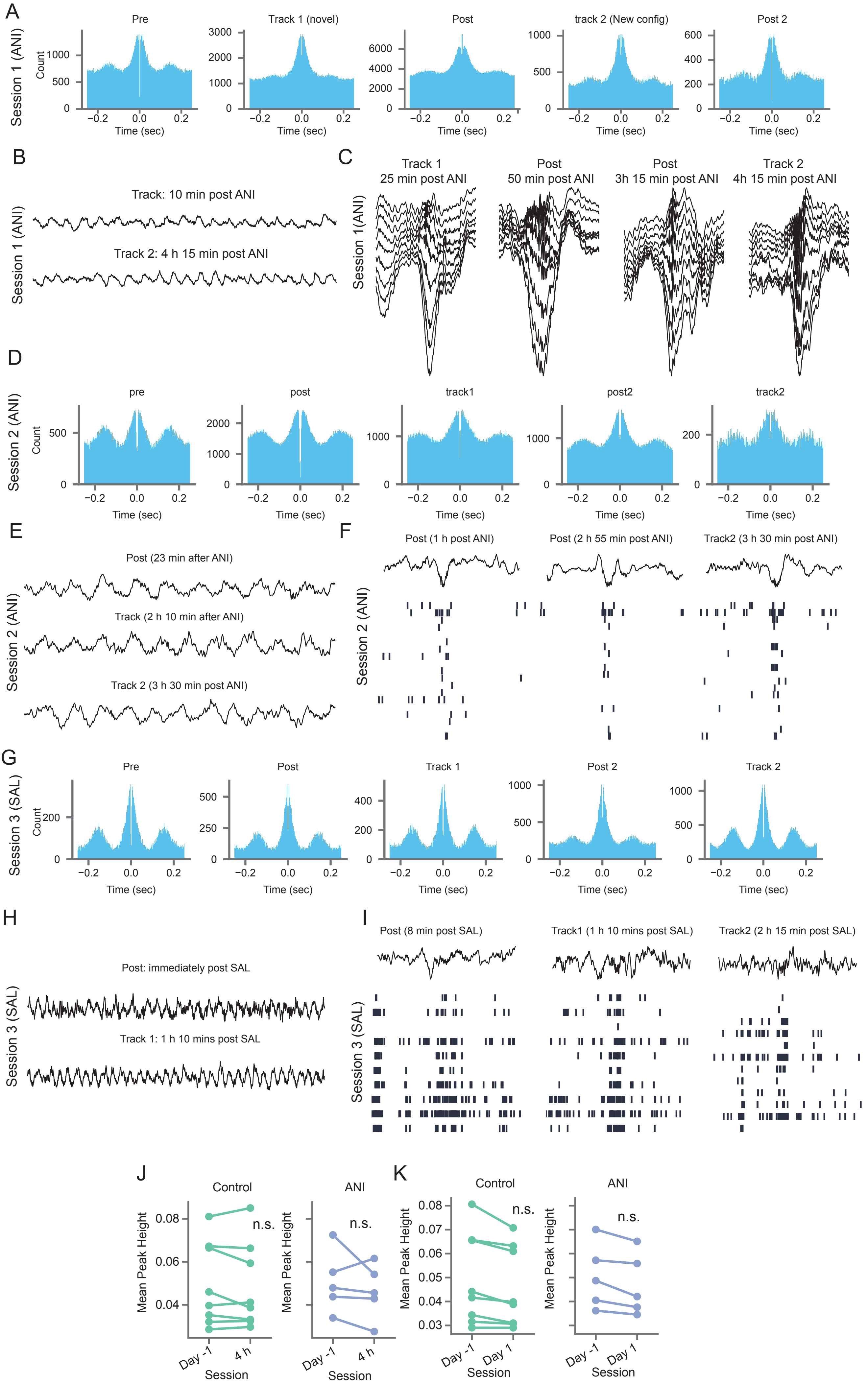


**Figure S5: Reduced activity following anisomycin administration is not an imaging artifact and does not result from global disruption of electrical neural activity in hippocampal neurons.**
**A-C)** Neural activity was tracked across ~5 hours before and after systemic administration of anisomycin in a rat. **A)** Cross correlograms for all single and multi-unit activity combined are shown from the pre epoch in a rest box (15 minutes), running on a novel track immediately following anisomycin injection (45 minutes), post epoch in the rest box (3.5 hours), running on a second novel track (45 minutes), and a second post epoch in the rest box (15 minutes). Clear modulation of firing at the theta timescale is observed. **B)** Example trace from electrode in pyramidal cell layer of CA1 showing theta activity 10 minutes and 4 hours 15 minutes post injection anisomycin injection across 9 channels of a linear probe spanning above to below the pyramidal cell layer. **C)** Example sharp wave ripple events occurring from 25 minutes to 4 hours 15 minutes post anisomycin injection. **D-F)** Same as A-C but for a different rat following systemic anisomycin injection. **G-I)** Same as D-F but the following day after systemic saline (control) injection demonstrating no lasting effects of anisomycin 24 hours after injection. F) Shows one trace from an electrode in the cell layer with a raster of spiking activity from all units recorded shown below the trace, demonstrating a population burst coincident with each sharp wave ripple. **J-K)** The signal-to-noise ratio of all mouse neurons captured using calcium imaging and active between sessions was tracked between sessions. **J)** Mean height of calcium transient peaks for all cells matched from day -1 to 4 hour session. $p>0.63$ both groups, two-sided t-test. (n=8 CTRL and 7 ANI, 1 additional CTRL mouse included whose video tracking behavioral data was corrupted and could therefore could not be classified as a Learner or Non-Learner). **K)** Same as J) but tracking cells from day -1 to day 1, $p>0.68$ both groups.

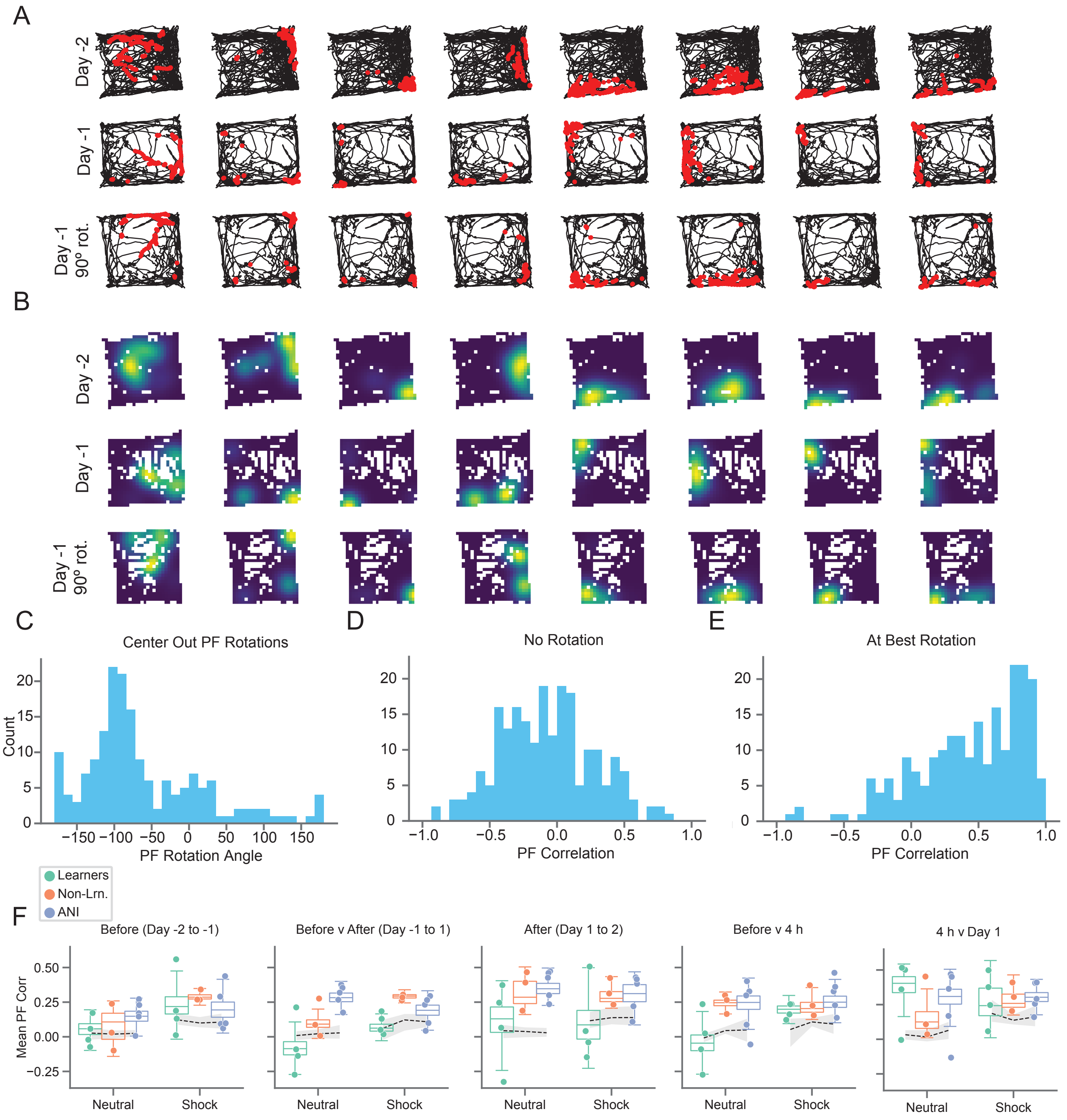


**Figure S6: Coherent Place Field Rotations Observed Between Sessions**

**A)** Example animal trajectories from neutral arena day -2 (top row) and day -1 (middle row) with calcium activity overlaid (red). Each column corresponds to one cell. Bottom row shows data rotated 90 degrees, demonstrating a coherent rotation of spatial activity for all neurons. **B)** Smoothed, occupancy normalized calcium event maps corresponding to data shown in A). **C)** The angle from the center of the arena to each cell's maximum intensity place field center was calculated for each session (center-out angle).
The distribution of center-out angles plotted, demonstrating a coherent rotation of place fields from Day -2 to Day -1 by 90 degrees. **D)** Place field correlations (smoothed event maps) between sessions indicate apparently low stability across days without considering rotations, giving the false impression that the place field map randomly reorganizes between sessions. **E)** High correlations were observed after considering a coherent 90 degree rotation between sessions, indicating that place fields retain the same relative structure but rotate together as a whole. **F)** Mean correlations for each mouse without considering rotations gives the impression of instability before/after shock and heightened remapping for all groups from before to after learning. Boxplots show population median and 1st/3rd quartiles (whiskers, 95% CI) estimated using hierarchical bootstrapping (HB) data with session means overlaid in dots. Dashed line and grey shading show mean and 95% CI of correlations calculated from shuffling cell identify 1000 times between sessions.

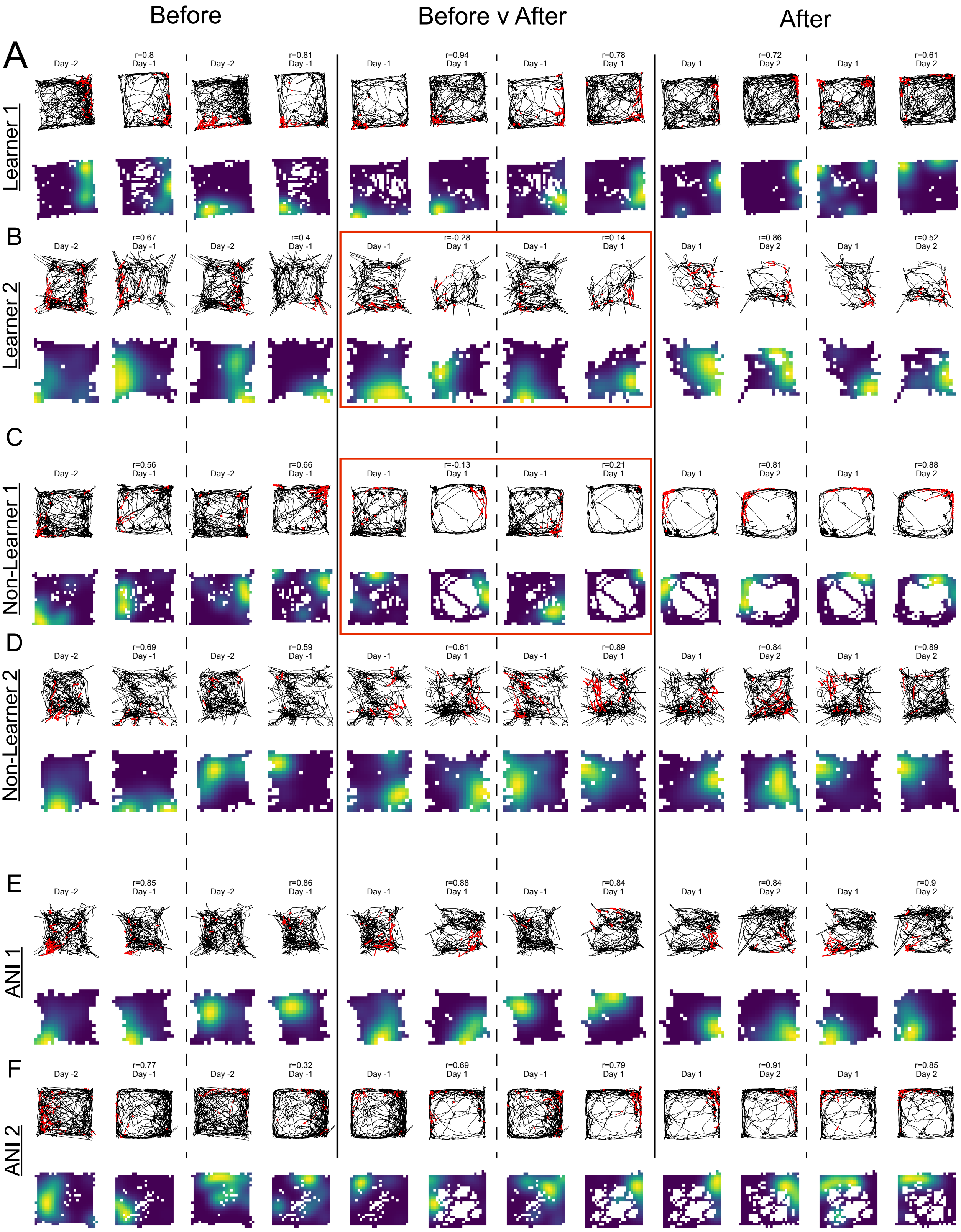


**Figure S7: Example stable and remapping place fields across sessions**

**A)** Example place field plots for Learner mouse in Neutral arena Before conditioning (Day -2 to Day -1), from Before to After conditioning (Day -1 to Day 1), and After consolidation (Day 1 to Day 2) demonstrating high stability in Neutral arena across all time points. (top row) Mouse trajectory in black with cell calcium activity overlaid in red. (bottom row) Smoothed, occupancy normalized calcium event spatial maps corresponding to raw trajectory and event data shown in top row, with warmer colors indicating high event rates and cool colors indicating low event rates. Spearman correlation value between event rate maps shown at top. Dashed lines denote separate different cells at each time point, and solid lines separate different comparison times. **B)** Same as A) but for different Learner in Shock arena demonstrating remapping. Red box shows two example remapping cells in the Shock arena. **C) and D)** Same and A) and B) but for two different Non-Learners with red box showing remapping cells in the Neutral Arena. **E) and F)** Same as A) and B) but for two different ANI group mice showing stable place fields between sessions at all time points.

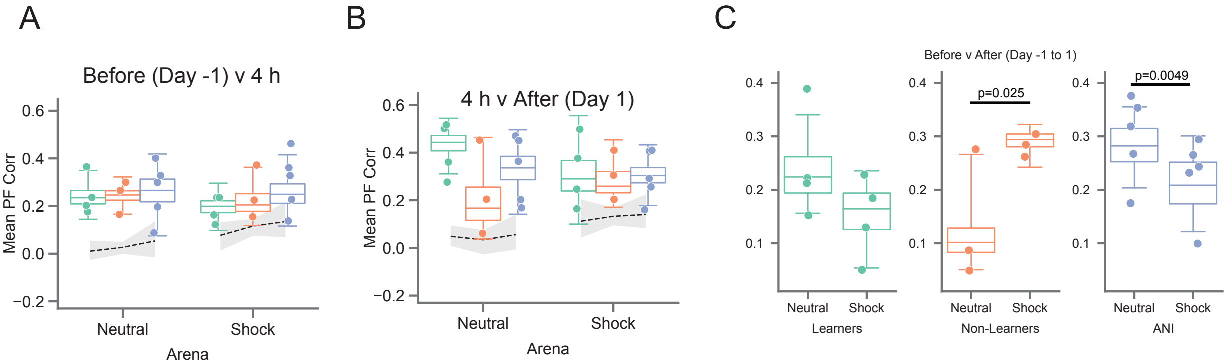


**Figure S8: Place field correlations with STM (4 hour) session**

**A)** Place field correlations for all mice combined for Day -1 vs 4 hour session. Boxplots show population median and 1st/3rd quartiles (whiskers, 95% CI) estimated using hierarchical bootstrapping (HB) data with session means overlaid in dots. Dashed line and grey shading show mean and 95% CI of correlations calculated from shuffling cell identity 1000 times between sessions. **B)** Same as A) but for 4 hour vs Day 1 session. All hierarchical bootstrap test comparisons are ns. **C)** Place field correlations for all mice in each group from before to after shock (Day -1 to Day 1). Boxplots show population median and 1st/3rd quartiles (whiskers, 95% CI) estimated using hierarchical bootstrapping (HB) data with session means overlaid in dots. Significant p-values calculated using a one-sided HB permutation test are shown directly on each panel.

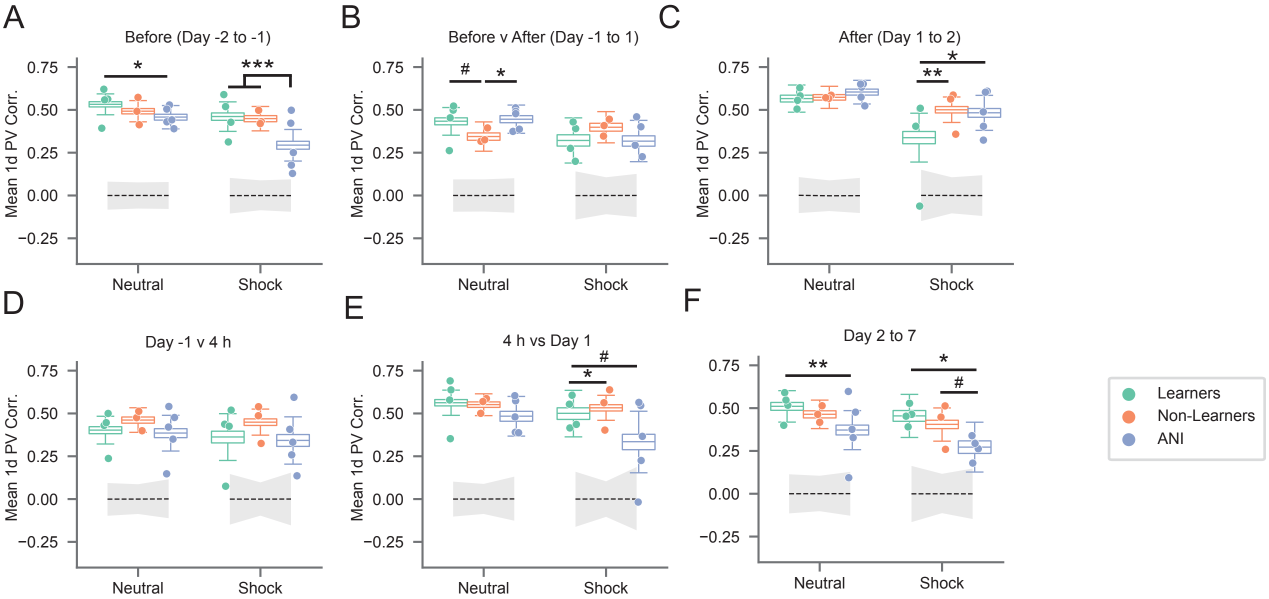


**Figure S8: Population Vector (PV) correlations.**
**A)-E)** 1D PV correlations between sessions including only cells active in BOTH sessions. **A)** Before (Day -2 vs -1), *p=0.042, ***p<0.0006. **B)** After (Day 1 vs 2) *p=0.034, #p=0.072. **C)** Before v After (Day -1 vs 1), *p<0.039, **p=0.0098**. D)** Day -1 vs 4 hr session. **E)** 4 hr session vs Day 1 *p=0.0049, #p=0.076. **F)** Day 2 vs Day 7, *p=0.0189, **p=0.0164, #p=0.064. Green = Learners, Orange = Non-Learners, Blue = ANI. Boxplots show population median and 1st/3rd quartiles (whiskers, 95% CI) estimated using hierarchical bootstrapping (HB) data with session means overlaid in dots. Dashed line and grey shading show mean and 95% CI of correlations calculated from shuffling cell identify 1000 times between sessions. Statistics: un-paired one-sided HB test after Bonferroni correction.

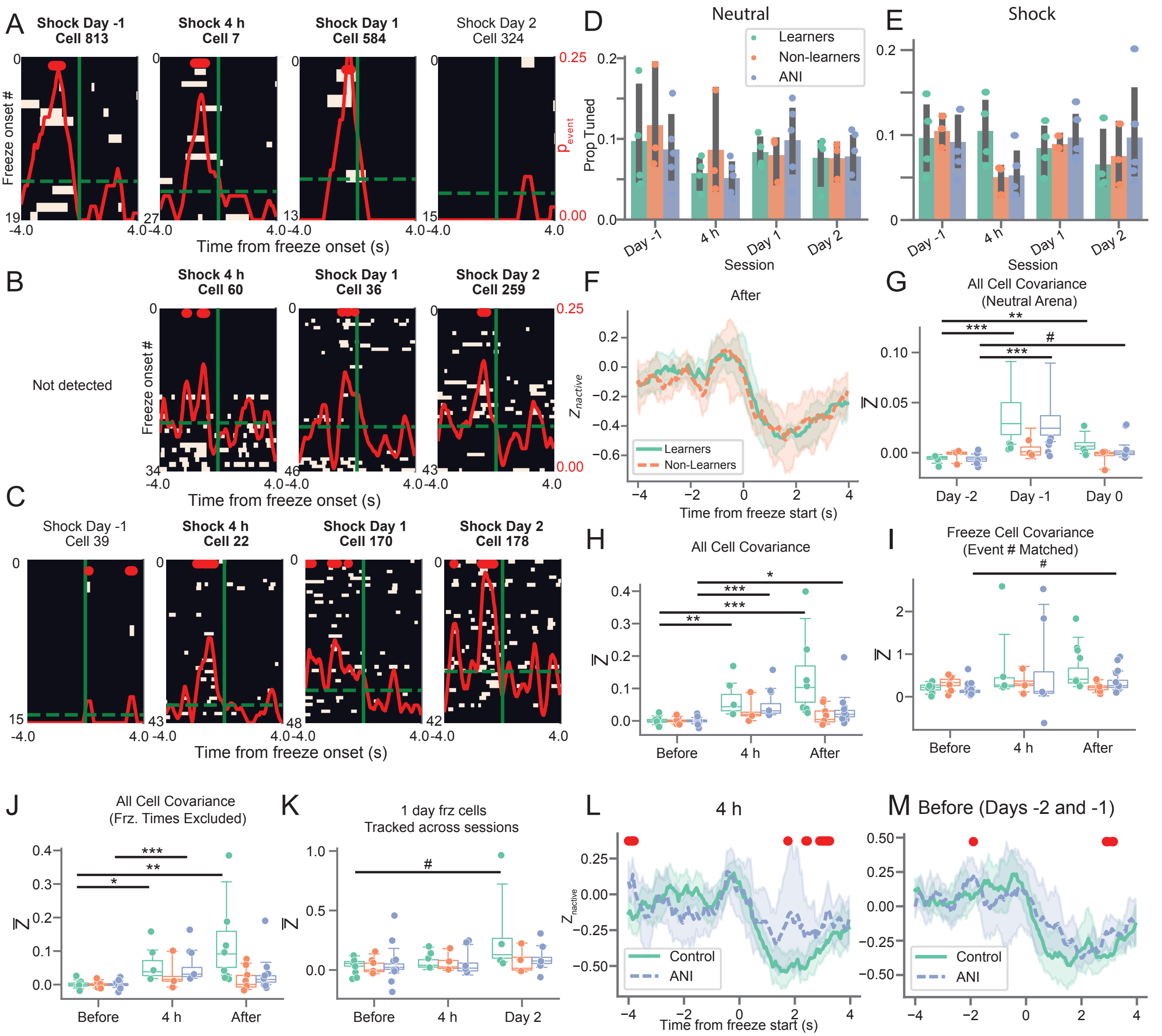


**Figure S10: Freeze-cell covariance increases are driven by Learners and not purely a result of less freezing in Non-Learners and the ANI group.**

**A)-C)** Example freeze-predicting cells tracked across sessions forward and backward in time from the day indicated in bold. Peri-event calcium activity rasters are centered on freeze onset time (solid green). Dashed green = baseline calcium event probability, red solid = peri-freeze calcium event probability, bins with p<0.01 (circular permutation test) noted with red bars at top. D/E corresponds to pink/blue cells shown in A-B. **A)** Example freeze-predicting cell from Non-Learner **B)-C)** Example freeze-predicting cells from two different Learners. **D)** Proportion freeze-predicting cells detected in Neutral arena across days. Bars=mean, line=std. **E)** Same as D) but for Shock arena. **F)** After learning (Days 1 and 2), z-scored population level calcium activity peaks between 0 and 2 seconds prior to freezing for both Learners and Non-Learners. Line/shading = mean +/- 95% CI. Red: bins with $p < 0.05$, independent t-test (two-sided, n = 4 Learners and 3 Non-Learners). **G)** Mean covariance of all cells in neutral arena prior to learning exhibit small changes, compare y-axis to Figure 3O and S6F, H.**p=0.0048, ***p<1e-8, #p=0.06. **H)** Same as G) but for all cells in shock arena. *p=0.026, **p=0.00052, ***p<2.5e-6. **I)** Same as G) but for freeze predictive cells only, peri-freeze times only, and after randomly downsampling the number of freeze events to match the average number observed during days -2 and -1. #p=0.056 **J)** Same as H) but excluding all peri-freeze times. *p=0.0018**p=1.2e-4, ***p=2.1e-5. **K)** Mean covariance of freeze-predicting cells detected on Day 1 tracked across time. #p=0.068. **L-M)** There are no significant differences between the proportion of active units (z-scored) peaks for Control animals compared to the ANI group between 0 and 2 seconds prior to freezing before conditioning (L) and at the 4 hour session (M). Line/shading = mean +/- 95% CI. Red: bins with $p < 0.05$, independent t-test (two-sided).

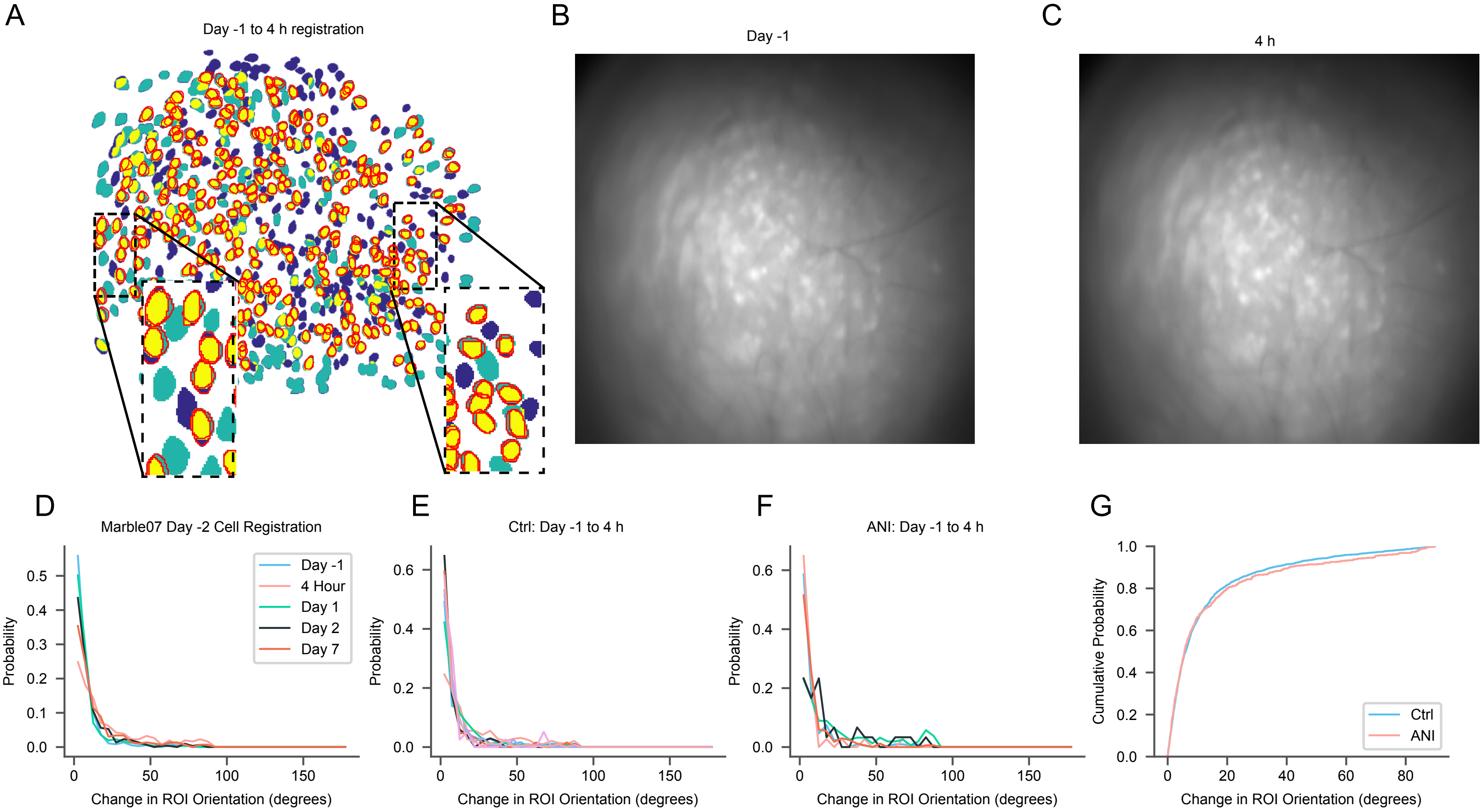


**Figure S11: Between Session Neuron Registration Metrics**
**A)** Example neuron registration between the Day -1 and 4 Hour session for one mouse in the shock arena depicting cell ROIs active during the Day -1 session only (blue), the 4 Hour session only (teal), and both the Day -1 and 4 Hour session (yellow) with ROIs which were successfully registered across sessions outlined in red. Insets show magnified ROIs and demonstrate that cells registered across days exhibit similar shape and orientation. **B)** and **C)** Minimum projections of the imaging movie from the sessions shown in A demonstrating high day-to-day stability, evidenced by clear alignment of landmarks (e.g. vasculature) between sessions. **D)** Change in ROI orientation for all neurons registered to the Day -2 session for the mouse shown in A). Note that the majority of registered neurons exhibit very small changes in orientation between sessions, even up to 9 days later (Day -2 to Day 7). **E)** and **F)** Similar to D) but for all mice in the CTRL and ANI groups separately for Day -1 to Day 4 (from before to after anisomycin administration). Note a similar distribution of ROI orientation changes, indicating that the observed acceleration of cell turnover following anisomycin administration is not due to poor neuron registration. **G)** Side-by-side comparison of all neuron ROI changes for each group shown as a cumulative density function.
